# Nuclear actin is a critical regulator of *Drosophila* female germline stem cell maintenance

**DOI:** 10.1101/2024.08.27.609996

**Authors:** Nicole M. Green, Danielle Talbot, Tina L. Tootle

**Affiliations:** Anatomy and Cell Biology, University of Iowa Carver College of Medicine, 51 Newton Rd, 1-500 BSB, Iowa City, IA 52242; Biology, Cornell College, 600 First Street SW, Mount Vernon, IA 52314; Biology, University of Iowa, 129 E. Jefferson St, 246 BB, Iowa City, IA 52242

**Keywords:** nuclear actin, oogenesis, *Drosophila*, germline stem cells, maintenance

## Abstract

Nuclear actin has been implicated in regulating cell fate, differentiation, and cellular reprogramming. However, its roles in development and tissue homeostasis remain largely unknown. Here we uncover the role of nuclear actin in regulating stemness using *Drosophila* ovarian germline stem cells (GSCs) as a model. We find that the localization and structure of nuclear actin is dynamic in the early germ cells. Nuclear actin recognized by anti-actin C4 is found in both the nucleoplasm and nucleolus of GSCs. The polymeric nucleoplasmic C4 pool is lost after the 2-cell stage, whereas the monomeric nucleolar pool persists to the 8-cell stage, suggesting that polymeric nuclear actin may contribute to stemness. To test this idea, we overexpressed nuclear targeted actin constructs to alter nuclear actin polymerization states in the GSCs and early germ cells. Increasing monomeric nuclear actin, but not polymerizable nuclear actin, causes GSC loss that ultimately results in germline loss. This GSC loss is rescued by simultaneous overexpression of monomeric and polymerizable nuclear actin. Together these data reveal that GSC maintenance requires polymeric nuclear actin. This polymeric nuclear actin likely plays numerous roles in the GSCs, as increasing monomeric nuclear actin disrupts nuclear architecture causing nucleolar hypertrophy, distortion of the nuclear lamina, and heterochromatin reorganization; all factors critical for GSC maintenance and function. These data provide the first evidence that nuclear actin, and in particular, its ability to polymerize, are critical for stem cell function and tissue homeostasis *in vivo*.

## INTRODUCTION

Actin was first reported to localize to the nucleus in the 1960s (Ohnishi *et al*. 1963; Ohnishi *et al*. 1964; Lane 1969). However, many dismissed these observations as artifacts for decades until the mechanisms controlling the nuclear import and export of actin were uncovered (Stuven *et al*. 2003; Dopie *et al*. 2012), and the invention of more advanced actin labeling tools (Melak *et al*. 2017; Kelpsch and Tootle 2018). It is now known that nuclear actin regulates a number of fundamental nuclear processes, including transcription, chromatin organization and remodeling, nuclear structure and integrity, nucleolar function and structure, and DNA damage repair (Melak *et al*. 2017; Kelpsch and Tootle 2018; Percipalle and Vartiainen 2019; Hyrskyluoto and Vartiainen 2020; Green *et al*. 2021; Kloc *et al*. 2021).

The ability to regulate essential nuclear functions makes nuclear actin a key factor in driving cell fate decisions. Nuclear actin is involved in regulating differentiation, deciding cell identity, and reprogramming cells to a pluripotent state (Misu *et al*. 2017; Xie *et al*. 2020; Green *et al*. 2021; Venit *et al*. 2021; Gunasekaran *et al*. 2022; Tomikawa and Miyamoto 2022). Nuclear actin mediates cell fate decisions by modulating the activities of factors such as RNA polymerase (RNAP) I, II, and III, specific transcription factors, histone deacetylase activity, and chromatin remodeling complexes. For example, in cancer, high levels of nuclear actin spur mammary epithelial cells to exit quiescence and assume a tumorous fate (Spencer *et al*. 2011; Fiore *et al*. 2017).

One way nuclear actin can influence cell fate is through the presence of specific nuclear actin structures. In the nucleus, actin can exist in many forms, including monomers, short polymers, and more extensive filamentous structures (Ulferts *et al*. 2021). Different forms of actin can be detected with labeling reagents specific to monomeric (e.g. DNase I; (Hitchcock 1980)) and polymeric forms (e.g. AC15; (Cruz and Moreno Diaz de la Espina 2009; Dopie *et al*. 2012; Wineland *et al*. 2018)). In the cytoplasm, actin functions are dependent upon actin structure; similarly, we see that nuclear actin structure is important for regulation of many nuclear functions (Kelpsch and Tootle 2018; KyherÖINEN AND VARTIAINEN 2020; Green *et al*. 2021; Ulferts *et al*. 2021). In particular, the advent of new actin labeling tools revealed that nuclear actin polymers appear at distinct and critical points in development (Grosse and Vartiainen 2013; Melak *et al*. 2017; Plessner and Grosse 2019). These findings suggests that nuclear actin polymerization is important for cellular events required at developmental transitions (Plessner *et al*. 2015; Baarlink *et al*. 2017; Misu *et al*. 2017; Okuno *et al*. 2020; Gunasekaran *et al*. 2022; Mahmood *et al*. 2022). Indeed, studies show that nuclear actin polymerization is required for establishing specific transcriptional programs, organizing chromatin structures, and differentiating into unique cell identities (Ferrai *et al*. 2009; Sen *et al*. 2015b; Le *et al*. 2016; Sen *et al*. 2017; Boraas *et al*. 2018; Sankaran *et al*. 2019b; Wei *et al*. 2020). Furthermore, somatic nuclear reprogramming in Xenopus oocytes requires nuclear actin polymers to drive expression of pluripotency genes (Miyamoto *et al*. 2011b). These observations highlight the importance of nuclear actin structure in cell fate, yet the mechanistic details of how nuclear actin structures are coordinated with cellular fate and drive specific nuclear functions is only beginning to be understood. Therefore, there is a critical need to learn more about the regulation of nuclear actin levels, as well as nuclear actin structures, to understand how nuclear actin contributes to cell fate and tissue development.

To examine nuclear actin’s role in cell fate in an *in vivo* context, we used a model stem cell population within the *Drosophila* ovary, the germline stem cells (GSCs). The *Drosophila* ovary is made up of 15-20 ovarioles of sequentially developing follicles (Giedt and Tootle 2023). GSCs give rise to all germline cells and reside in a niche at the anterior tip of each ovariole in a structure known as the germarium (Fig. 1A). Each GSC nucleus is large, comprising the majority of the cell, has a specific nuclear lamina composition and exhibits a large, round nucleolus surrounded by the nucleoplasm (Fig. 1B; (Duan *et al*. 2020)).

**Figure 1.**
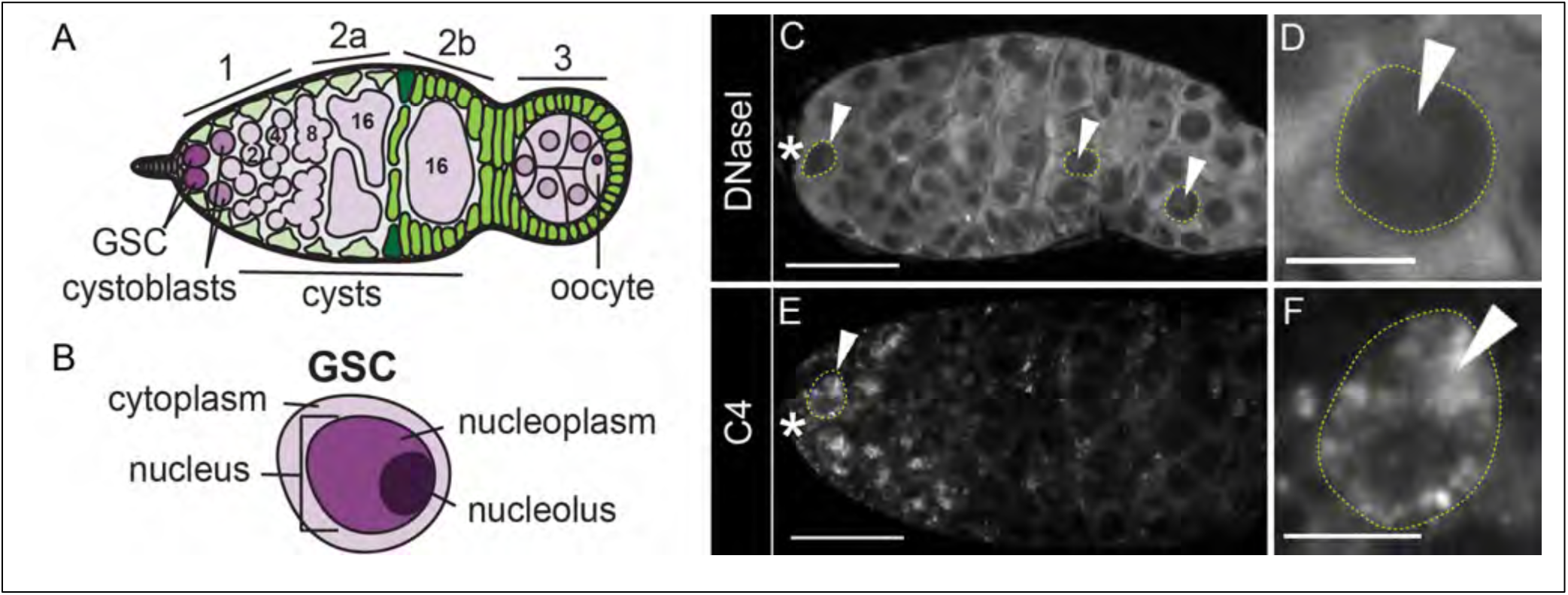
Nuclear actin pools in the *Drosophila* germarium. (A-B) Schematic of a germarium with regions designated (A) and a germline stem cell (GSC, B); germline is shown in purple and somatic cells in green. (C-F) Maximum projections of 2-4 confocal slices of wild-type (*yw*) germaria stained with nuclear actin reagents: fluorescently conjugated DNase1 (C-D) and C4 anti-actin antibody (E-F). Asterisks designate the anterior tip and GSC niche in each germarium. Yellow dotted lines outline example germline nuclei with nuclear actin staining; D, F are cropped images of a single GSC. Arrowheads indicate nuclear actin pools in each germline cell. Scale bars = 15 μm in C, E; 5 μm in D, F. There are two major cell populations that make up the *Drosophila* ovary (A): 1) germline cells (purple), which give rise to the developing oocyte and are derived from the germline stem cells (dark purple; GSCs), and 2) the somatic cells (green) which surround the germline and provide important signaling cues and structural support; follicle stem cells are shown in dark green. In Region 1 of the germarium (A), two to three GSCs reside in an anterior niche. A GSC (dark purple) will asymmetrically divide to produce two daughter cells–a GSC which will be maintained in the niche to renew the GSC pool, and a daughter cell or cystoblast (CB; purple) which goes on to divide mitotically to produce interconnected germline cysts. Mitotic division of CBs produce germline cysts of 2-, 4-, 8-, and 16-cells which become progressively more differentiated as they move from region 1 through region 2a. In region 2b, 16 cell cysts (light purple) become encapsulated by somatic cells (green) and move into region 3 where oocyte specification is completed and a number of specialized developmental processes in the germline begin. The subcellular features of GSCs include a minimal amount of cytoplasm (light purple), and two major nuclear compartments: 1) the phase separated, membrane-less nucleolus (dark purple) and 2) the nucleoplasm (purple), which comprises the remaining portion of the nucleus (B). Nuclear actin pools change with germline differentiation. DNase1 stains monomeric actin in the cytoplasm and nucleus of every germline cell (C, arrowheads), including in the GSCs (D). Whereas anti-actin C4 antibody (E-F) stains a more restricted population of nuclear actin in the anterior tip of germaria where GSCs and less differentiated germline cysts are located (E). High levels of C4 nuclear actin are present in GSCs (E-F, arrowheads) and C4 nuclear staining decreases between region 1 and region 2a and is lost as germline cells move through regions 2a and 2b.

*Drosophila* oogenesis has proved to be an exceptional model for studying cytoplasmic actin dynamics (Hudson and Cooley 2002) and more recently has been used to investigate actin’s role in the nucleus (Kelpsch *et al*. 2016; Sokolova *et al*. 2018; Wineland *et al*. 2018; BorkÚTI *et al*. 2022; Talbot *et al*. 2023). Monomeric actin localizes to the nucleus of all cells in the *Drosophila* ovary (Wineland *et al*. 2018). In later stages, nuclear actin regulates nucleolar structure and functions, such as ribosomal RNA (rRNA) transcription (Talbot *et al*. 2023). Previously, we screened available actin labeling tools and found three that stained actin populations in the ovary–fluorescently conjugated DNase I, anti-actin AC15, and anti-actin C4 (Kelpsch *et al*. 2016; Wineland *et al*. 2018). In this study, we use these reagents to examine distinct subcellular actin populations or ‘pools’ in the *Drosophila* germarium. As seen in prior work (Wineland *et al*. 2018), DNase I specifically labels monomeric actin and, in addition to staining the cytoplasm, it labels a nuclear structure in every cell in the germarium (Fig. 1C-D). Surprisingly, within the germarium, C4 labeled actin is present in GSCs and C4 actin is lost as cells become more differentiated (Fig. 1E-F). These data led us to question what roles specific forms of nuclear actin might play in GSCs and whether nuclear actin is important in maintaining stemness.

Here we present our novel findings that nuclear actin is present in *Drosophila* GSCs and is required to maintain germline stem cells. We identify a unique pool of nuclear actin labeled by the C4 antibody which dynamically localizes in early germ cells. In GSCs, there is both a nucleoplasmic C4 actin pool consisting of structured, polymeric actin, and a nucleolar pool, which is predominantly monomeric. The loss of the polymeric, nucleoplasmic pool early in differentiation led us to hypothesize that nuclear actin polymerization is required for regulating stemness. Indeed, overexpression of nuclear actin monomers that cannot polymerize causes progressive germline loss, suggesting that nuclear actin polymerization is required for maintaining GSCs. Preventing actin polymerization in GSCs results in dramatic changes to nuclear organization including nuclear and nucleolar hypertrophy, increased heterochromatin, and distortions of the nuclear lamina. Consistent with our hypothesis, restoring nuclear actin polymerization in GSCs rescues the progressive germline loss observed in ovaries with monomeric nuclear actin overexpression. These data lead to the model that regulation of nuclear actin polymerization controls nuclear architecture, and related nuclear functions, to maintain GSC stemness in oogenesis. Based on the high conservation of actin across all organisms and the essential nature of nuclear processes in maintaining cell fate, we anticipate that this model may be observed in a broad array of biological systems and their stem cell populations.

## METHODS

See Supplementary Table S1 for detailed information on the reagents used in these studies.

### Fly Husbandry & Stock Information

Fly stocks were maintained at 21°C on a Blooming Drosophila Stock Center (BDSC) cornmeal medium. Prior to dissection, flies were fed with yeast paste daily for 4 days (unless otherwise indicated). Wild-type controls were *yw* flies unless otherwise specified; for GAL4-UAS experiments both GAL4 only and UAS-construct only controls were performed. The following stocks were obtained from the Bloomington Stock Center (Bloomington, IN): *yw* (BDSC1495), *Tmod-GFP* (BDSC50861), and *matα-*GAL4 (third chromosome; BDSC7063). The following stocks were generous gifts: *UASp-NLS-FLAG-Act5C* (NLS-Act-WT) and *UASp-NLS-FLAG-Act5C(G13R)* (NLS-Act-G13R) obtained from Maria Vartiainen (University of Helsinki) and generated by Peter Vilmos (Biological Research Center of the Hungarian Academy of Sciences), *nos(VP16)-GAL4* stock from Pamela Geyer (University of Iowa), and *vasa-*GAL4*, UAS-GFP* from Michael Buszczak (University of Texas Southwestern). Overexpression of *NLS-FLAG-Act* constructs was performed by crossing to *nos-*GAL4, *vasa*-GAL4 or *matα-*GAL4, maintaining crosses at 18°C through early gonadal development with a shift to 25°C at eclosion for 2-14 days (see individual experiments for details).

### Immunofluorescence

*Drosophila* ovaries were dissected in Grace’s insect medium (Lonza, Walkersville, MD), and fixed for 15 min on a nutator at room temperature in 4% paraformaldehyde in Grace’s insect medium. Samples were washed and permeabilized using Triton antibody wash (1x phosphate-buffered saline (PBS), 0.3% Triton X-100, and 0.1% bovine serum albumin (BSA; Sigma-Aldrich, St. Louis, MO) six times for 10 min each at room temperature. Primary antibodies were incubated for a minimum of 24 hr at 4°C on a nutator. The following primary antibodies were obtained from the Developmental Studies Hybridoma Bank (DSHB) developed under the auspices of the National Institute of Child Health and Human Development and maintained by the Department of Biology, University of Iowa (Iowa City, IA): rat anti-Vasa, 1:100 (RRID:AB_760351); mouse anti-Hts, 1:50 (RRID: AB_528070, 1B1); and mouse anti-LaminDm0, 1:200 (RRID:AB_528338, ADL84.12). The following primary antibodies were also used: mouse anti-actin C4, 1:50 (RRID: AB_2223041, EMB Millipore, Billerica, MA); rabbit anti-Fibrillarin, 1:250 (RRID:AB_2105785, Abcam, Cambridge, MA); goat anti-Otefin/Emerin, 1:500 ((Barton *et al*. 2014); gift from P. Geyer); mouse anti-FLAG M2, 1:300 (RRID:AB_262044, Sigma-Aldrich, St. Louis, MO); rabbit anti-GFP, preabsorbed at 1:20 and used 1:100 (RRID: AB_10013661, Torrey Pines Biolabs, Houston, TX); and rabbit anti-H3K9me3, 1:1000 (RRID:AB_2532132, Active Motif, Carlsbad, CA). Ovaries were washed with Triton antibody wash six times for 10 min each and then incubated in secondary antibodies for 12-48 hrs at 4°C on nutator. The following secondary antibodies (Life Technologies, Grand Island, NY) were used at 1:400: AF633::goat anti-mouse (RRID: AB_2535718), AF488::goat anti-mouse (RRID: AB_2534069), AF568::goat anti-rabbit (RRID: AB_10563566), AF488::goat anti-rat (AB_2534074), AF568::donkey anti-goat (RRID: AB_2534104), AF488::donkey anti-mouse (RRID: AB_141607), AF647::donkey anti-rabbit (RRID: AB_2536183). When used, AF488-DNase I, 1:500 (Invitrogen, Waltham, MA) or AlexaFluor568-conjugated Phalloidin, 1:500 (ThermoFisher Scientific, Waltham, MA) was added in both primary and secondary antibody incubations. After six additional washes in Triton antibody wash, ovaries were stained with 4’,6-diamidino-2-phenylindole (DAPI, 5 mg/mL, RRID: AB_2629482) diluted 1:5,000 in PBS for 10 min at room temperature, followed by a final wash in PBS. Ovaries were teased apart on slides and mounted in VectaShield mounting medium (RRID: AB_2336789, Vector Laboratories, Burlingame, CA). For tile scans, ovaries were left intact throughout the staining and gently teased directly on slide to prevent loss of ovarioles during quantification. All stainings were performed a minimum of three times.

### Image acquisition and processing

Images of fixed *Drosophila* follicles were obtained using Zen software on either a Zeiss 700 or Zeiss 880 mounted on a Zeiss Axio Observer.Z1 using Plan-Apochromat 20x/.8 working distance (WD)=0.55 M27 or Plan-Apochromat 63×1.4 Oil DIC f/ELYRA objective (Carl Zeiss, Thornwood, NY). Tile scans of whole ovaries were collected using 3×3 tiles at 20x. Tile scans were stitched with a 10% overlap and assembled into maximum projections using Zen Black software (RRID: SCR_018163, Carl Zeiss, Thornwood, NY). For germarium images, single image slices or maximum projections (2-4 confocal slices), merged images, rotation, and cropping were performed using FIJI software (RRID: SCR_002285; (Abramoff *et al*. 2004)). To aid in visualization, panels of the same figure were uniformly brightened using Photoshop (RRID: SCR_014199, Adobe, San Jose, CA) or Illustrator (RRID: SCR_010279, Adobe, San Jose, CA) as described in each figure legend.

### Western Blot

At eclosion, control and NLS-actin overexpression females were shifted from 18°C to 25°C to initiate construct expression. To minimize ovary size differences caused by germline loss, construct expression was induced for only 24 hours which precedes any observed germline loss. Whole ovaries were then dissected in room temperature Grace’s insect medium transferred to 50 µl 1x Laemmli buffer in a 1.5 mL microcentrifuge tube and homogenized by plastic pestle. Lysates were boiled at 100°C for 10 minutes and centrifuged at 15,000 x g to pellet cell debris. Samples were run on a 10% SDS-PAGE gel with the Precision Plus Protein All Blue Prestained Protein Standard (BioRad Laboratories, Hercules, CA) and transferred to a nitrocellulose membrane (Cytiva, Wilmington, DE). Variation in sample preparation was normalized by diluting samples based on germline content determine by western blotting for Vasa. Blots were blocked in a 5% milk primary antibody solution for 30 minutes (1x Tris-buffered saline (1X TBS) with 0.1% Tween-20). The following primary antibodies were used: rabbit anti-FLAG, 1:2000 (RRID:AB_262044, Sigma-Aldrich, St. Louis, MO) and rat anti-vasa, 1:100 (RRID:AB_760351, DSHB). Following primary antibody incubation, blots were washed three times with 1x TBS and one time with 1x TBS with 0.1% Tween-20 for 10 minutes each. The following secondary antibodies were used at 1:5000 in 5% milk in 1x TBS with 0.1% Tween-20: Peroxidase-AffiniPure Goat Anti-Rabbit IgG (H+L) (RRID: AB_2313567, Jackson ImmunoResearch Laboratories, West Grove, PA), and Peroxidase-AffiniPure Goat Anti-Rat IgG (H+L) (RRID: AB_2338128, Jackson ImmunoResearch Laboratories, West Grove, PA). Following secondary antibody incubation, blots were washed three times with 1x TBS and two times with 1x TBS with 0.1% Tween-20 for 10 minutes each. Blots were developed with SuperSignal West Pico Chemiluminescent Substrate (Thermo Scientific, Waltham, MA) and imaged using the Amersham Imager 600 (GE Healthcare Life Sciences, Chicago, IL). A minimum of three independent experiments were performed for each Western blotting experiment. Quantification of the western blot was performed by obtaining the raw intensity values from the Amersham Imager 600. The flag raw intensity value was then divided by the vasa raw intensity value to get an experimental to control ratio. All ratios were then divided by the average of the *nos-*GAL4/+ ratios to determine the relative amount of construct expression. Full blots are provided in Figure S2.

### Quantification of ovarian phenotypes

Flies were raised at 18°C until eclosion, collected in a fresh vial, and then shifted to 25°C for timepoints of 2, 4, or 14 days. To quantify germline loss, we analyzed tile scans of whole ovaries by cross referencing the Vasa channel (germline stain) with the DAPI channel (stains both germline and somatic cells). Images of ovary tile scans were blinded and each germarium was classified as one of the following phenotypic classes: 1) normal, where a range of germline cysts were present; 2) emptying, which included any germarium that had lost some germline cysts; 3) empty, where all germline cells (Vasa positive) were missing and only somatic cells remained (labeled with DAPI only). When necessary, high magnification images of DAPI channel were used to confirm ovarioles that were emptying or completely empty (DAPI positive only). The data from a minimum of three trials were compiled, pooled, and phenotypic percentages were calculated at each timepoint in Excel (RRID: SCR_016137, Microsoft, Redmond, WA). Data was graphed as a stacked bar graph using GraphPad Prism 9 (RRID: SCR_002798, GraphPad Software, San Diego, CA).

## RESULTS

### A dynamic pool of nuclear actin is present in the early female germline

Previously, we found that an actin antibody, anti-actin C4, labeled nuclei in the *Drosophila* germarium (Wineland *et al*. 2018). Unlike DNase I positive nuclear actin, which is found in all nuclei (somatic and germline), C4 nuclear staining is present in a subset of cells in the germarium (Fig 1E). Here we focus solely on the germline cells. The C4 antibody stains actin in the nuclei of GSCs as well as early germline cysts, but this staining decrease as cysts become more differentiated (Fig. 1E-F). As previously described, nucleolar C4 actin is also observed in a subset of the nurse cells in region 3 of the germarium and in early stages of follicle development (Kelpsch *et al*. 2016; Wineland *et al*. 2018). The dynamic C4 staining pattern within the germarium suggested nuclear actin could be an important factor for regulating GSCs and maintaining a stem cell fate.

Here we characterize the cell-specific C4 staining pattern in the GSCs and find C4 labels actin in distinct nuclear regions (Fig. 2). Nuclei contain two major sub-compartments: 1) the nucleolus, a phase-separated functional compartment (Fig. 2A-B’, Fibrillarin), and 2) the remaining nucleoplasm which is surrounded by the nuclear lamina (Fig. 2A-B”, Emerin). To define the subcellular distribution of nuclear actin in the germarium, we co-labeled wild-type ovaries with the C4 antibody and several key nuclear landmarks – the nuclear lamina, chromatin, and the nucleolus. GSCs exhibit C4 nuclear actin in both the nucleoli (Fig. 2, yellow solid line) and nucleoplasm (Fig. 2, yellow dotted line). Within the nucleoplasm, C4 staining is irregular and lacks uniformity throughout the compartment (Fig. 2B-B’; C-D’). Because a major feature of the nucleoplasm is chromatin, we co-stained germaria to determine where C4 actin is found in relation to organized chromatin. C4 staining does not strongly overlap with trimethylated histone 3 lysine 9 (H3K9me3) marks on heterochromatin (Fig. 2C-C”, D-D’’) nor does it co-localize with regions of bright DAPI staining which broadly labels chromatin (Fig. 2, C’’’-D’’’).

**Figure 2.**
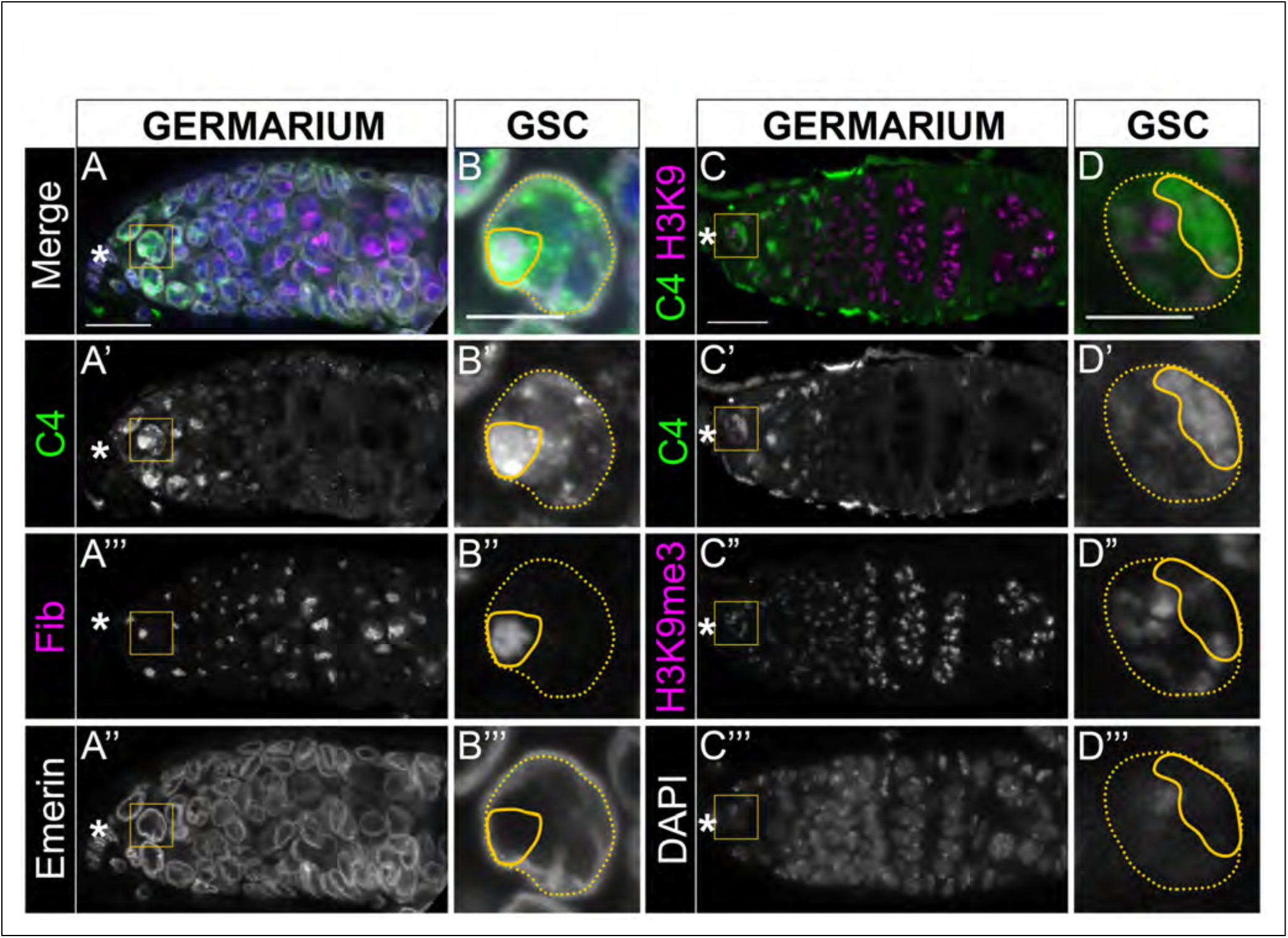
C4 nuclear actin pools are present in the nucleoplasm and the nucleolus of the early germ cells. (A-D’’’) Maximum projections of 2-5 confocal slices of germaria (A-A’’’, C-C’’’) and zoomed in images of GSCs in the yellow boxed regions (B-B’’’, D-D’’’’) of wild-type (*yw*) 4 day old flies stained with anti-actin C4 (green in merge) and various markers of nuclear organization including DAPI (blue in merge), Emerin (nuclear lamin associated protein, white in merge), Fibrillarin (Fib, nucleolar marker, magenta in merge), and H3K9me3 (trimethylation of lysine 9 on histone H3; heterochromatin marker, magenta in merge). Asterisks designate the anterior tip and GSC niche in each germarium. Note that the DAPI channel is presented only as a separate channel in C’’’ and D’’’ and is not included in the merge for clarity. Scale bars = 15 μm in A-A’’’, C-C’’’; 5 μm in B-B’’’, D-D’’’. C4 positive nuclear actin is present in two populations within the GSC nuclei –the smaller ‘nucleolar C4’ pool (A-B’ and C-D’, outlined by solid yellow line in B-B’ and D-D’) that overlaps with Fibrillarin (A-A’’’, B-B’’’) and the larger ‘nucleoplasmic C4’ pool (A-B’ and C-D’, outlined by yellow dashed line in B-B’ and D-D’) present throughout the rest of the nucleus as marked by either the nuclear lamina associated protein, Emerin (A, A’’ and B, B’’) or DAPI (C’’’ and D’’’). In the nucleoplasm, C4 staining is found in both heterochromatic and non-heterochromatic regions (C-D”’).

We then asked how the C4 nuclear actin pools change during germline cyst differentiation. Using the morphology of the fusome, an actin-rich structure, we staged the germline cells (Fig. 3A-D’’’, F-actin). GSCs are proximal to the cap cells and contain a special round fusome called a spectrosome (Fig. 3A’’’). As GSCs divide, the daughter cell called the cystoblast (CB) moves away from the niche and retains a rounded spectrosome structure (Fig. 3B’’’). Germline cysts of multiple cells (2-, 4-, 8-, 16-cell cysts) contain branched fusomes that exhibit a corresponding number of branches to connect all the cells within the cyst (Fig. 3B’’’-D’’’). In each cyst stage, we use DAPI (not shown) and Fibrillarin to distinguish the nucleoplasmic and nucleolar pools, respectively. The nucleoplasmic pool of C4 actin is present in GSCs through 2-cell cysts (Fig. 3A-B’). Nucleolar C4 actin persists longer and is present in the GSCs through 8-cell cysts (Fig. 3A-C”). Neither the nucleoplasmic nor nucleolar C4 nuclear actin is present in 16-cell cysts (Fig. 3D-D”). To confirm these results, we also used a Tropomodulin-GFP (Tmod-GFP) protein trap to label fusomes and observe the same developmental pattern of nucleoplasmic and nucleolar C4 staining (Fig. S1). These data reveal there are two distinct C4 pools in the early germline: an early nucleoplasmic C4 pool present in GSCs and less differentiated germline cysts, and a more persistent nucleolar C4 pool which is present in GSCs until the transition between the 8- and 16-cell cyst stage (Fig. 3E).

**Figure 3.**
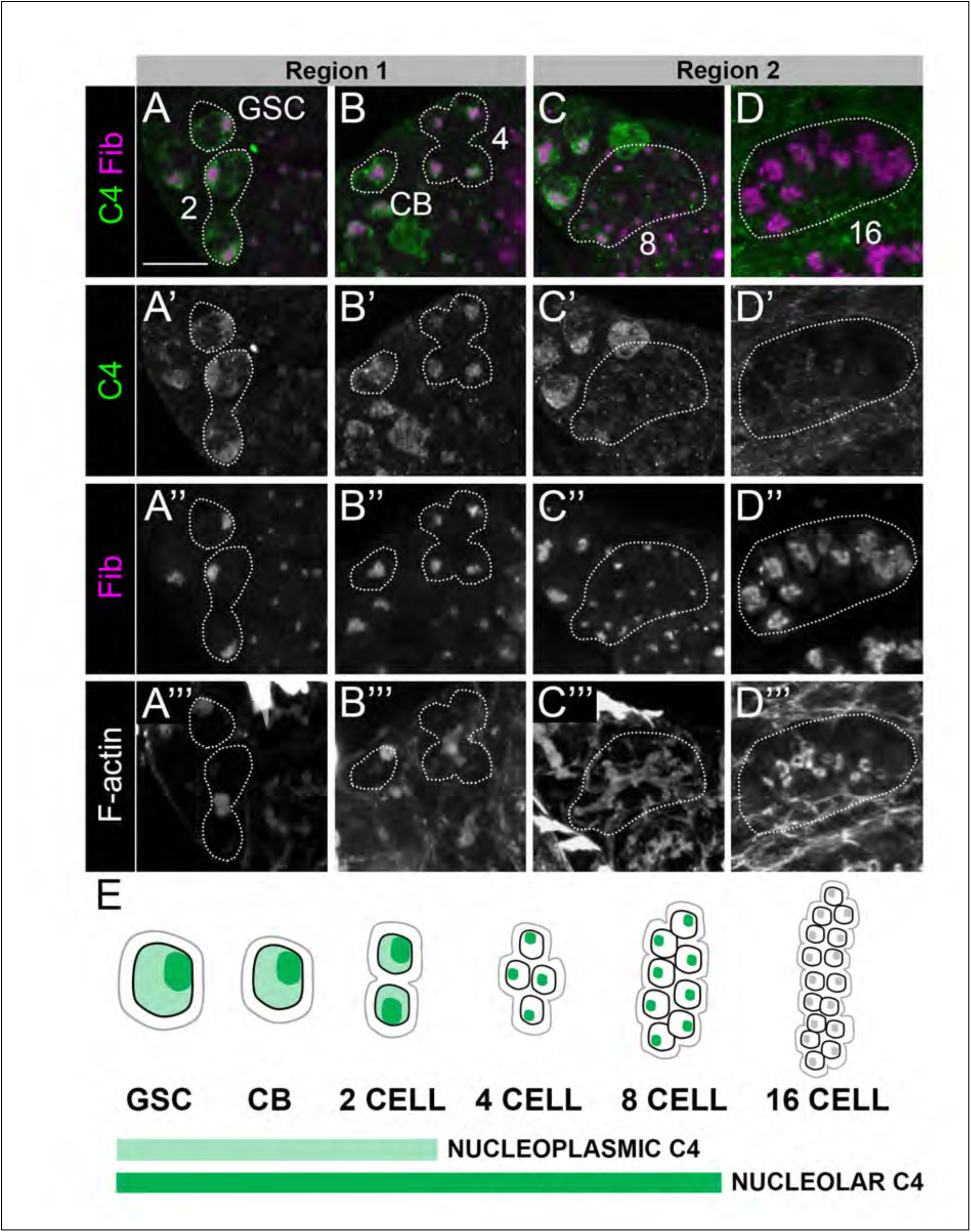
C4 nuclear actin is present in GSCs and decreases as germline cysts become more differentiated. (A-D’’’) Maximum projections of 5-10 confocal slices of wild-type (*yw*) germaria from 4 day old flies cropped to feature each stage of germline development stained with anti-actin C4 (green in merge, A-D and A’-D’), Fibrillarin to mark the nucleolus (Fib, magenta in merge, A-D and A”-D”), and F-actin (phalloidin) to mark the fusome (A’’’-D’’’). Note that the F-actin is presented only as a separate channel and is not included in the merged images for clarity. Germline cells and cysts were staged using fusome morphology and GSCs/cysts are outlined with white dashed line in all images: GSC=germline stem cell, CB=cystoblast, 2=2-cell cyst, 4=4-cell cyst, 8=8-cell cyst, 16=16-cell cyst (encapsulated cyst in region 2b pictured). Black boxes were added behind A’’’ and C’’’ to make letters visible on images. Scale bar = 10 μm. (E) Schematic summarizing the presence of nucleoplasmic C4 (light green) and nucleolar C4 (green) in germline nuclei across the germarium. Light gray line indicates cell membrane and the black line outlines the nucleus. Light gray circle within each nucleus indicate a nucleolus absent of C4 staining. The nucleoplasmic C4 pool is present in GSCs, CBs, and 2-cell cysts (A-B’’’, E), but is absent in the rest of germline cysts (B-D’’’, E). The nucleolar pool is present in GSCs through 8-cell cysts (A-C’’’, E), but is absent in 16-cell cysts (D-D’’’, E).

What is different about the two pools of C4 nuclear actin in the early germ cells? Previously, we found that both C4 and DNase I, which labels monomeric actin in all cells (Hitchcock 1980), colocalize to the early germ cell nucleoli (Wineland *et al*. 2018). Therefore, we think the C4 nucleolar staining corresponds to a monomeric nuclear actin pool (Fig. 3E). However, unlike DNase I, C4 does not label nucleolar actin in every cell in the germarium (Fig. 1C, E), indicating C4 stains a unique and likely modified form of monomeric actin in specific germline cells. The nucleoplasmic C4 staining does not overlap with DNase I. This finding leads us to think that nucleoplasmic C4 labels a polymeric form of actin (Wineland *et al*. 2018). As the nucleoplasmic C4 actin is restricted to the GSCs and early germline cysts (Fig. 3E), we hypothesize that this polymeric actin is a key determinant in deciding between a stem-like versus a more differentiated cell fate.

### Nuclear actin is required for GSC maintenance and germline development

The nucleoplasmic, polymeric C4 nuclear actin exhibits an inverse correlation with differentiation state (Fig. 3E), raising the possibility that polymeric nuclear actin is an important factor for regulating GSC maintenance. To test this possibility, we asked if altering the levels of polymeric and/or monomeric nuclear actin in the early germline disrupts GSC maintenance processes required to prevent germline loss. However, manipulating an essential molecule such as actin is technically challenging. The same actin found in the nucleus is used to make the actin cytoskeleton. Thus, knocking down actin is not a viable option. Another common mechanism of reducing nuclear actin is to impair it import. However, it has recently been shown that multiple mechanisms control the nuclear localization of actin in *Drosophila* (BORKÚTI *et al*. 2022). Therefore, we decided to increase nuclear actin in GSCs. Specifically, we targeted the different nuclear actin pools by overexpressing actin engineered with a nuclear localization sequence (NLS) and tagged with FLAG (NLS-FLAG-Act) using the UAS/GAL4 system. The Act5C isoform was selected because this isoform is expressed in all tissues consistently throughout development (BORKÚTI *et al*. 2022) and it is the main nuclear actin during *Drosophila* oogenesis (Spracklen *et al*. 2014); a FLAG tag was added for actin visualization.

We overexpressed two different UAS-NLS-FLAG-Actin constructs in the early germline using *nos*-GAL4: NLS-Act-WT and NLS-Act-G13R. NLS-Act-WT encodes a nuclear targeted actin that is able to incorporate into elongating nuclear actin structures (i.e. ‘polymerizable nuclear actin’), undergo post-translational modifications, and be exported from the nucleus like endogenous actin. Therefore, NLS-Act-WT overexpression is expected to increase both the monomeric and polymeric nuclear actin pools. NLS-Act-G13R encodes a nuclear targeted mutant actin that cannot polymerize (Posern *et al*. 2002), but can still be modified and exported from the nucleus. Thus, NLS-Act-G13R increases monomer nuclear actin, and likely impairs nuclear actin polymerization. When polymerizable NLS-Act-WT is overexpressed in the germline (Fig 4B-B’, E-E’), anti-FLAG staining reveals a nuclear haze (open arrowhead) and F-actin labeling at the cell cortex throughout the germarium (closed arrowhead). In contrast, overexpression of NLS-Act-G13R (Fig. 4C-C’, F-F’) yields no obvious structured actin in the germarium or later stages (closed arrowhead), but does result in a strong haze of FLAG-actin within germ cell nuclei (open arrowhead).

**Figure 4.**
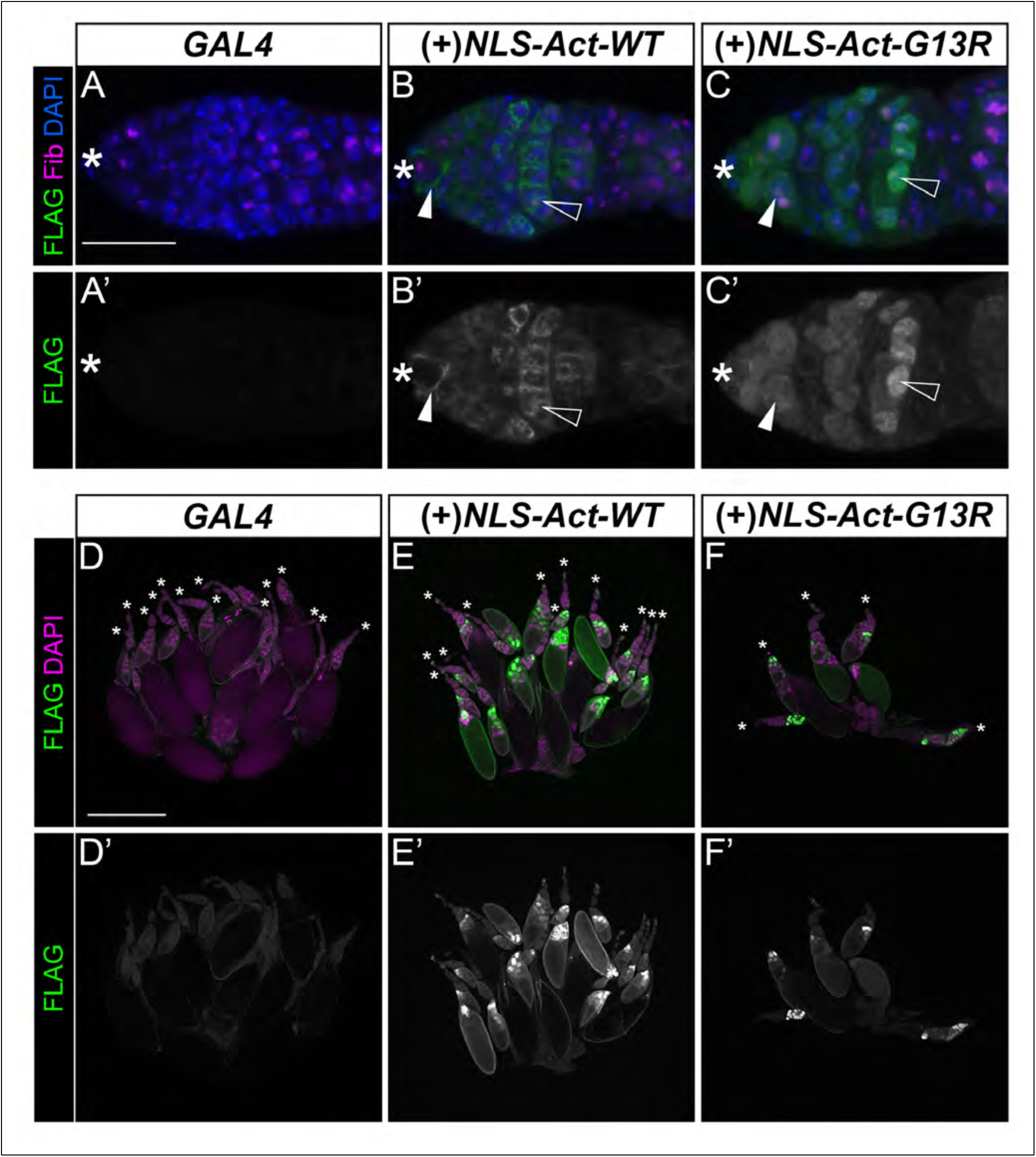
Germline overexpression of NLS-Act-G13R prevents nuclear actin polymerization and disrupts oogenesis. (A-C’) Maximum projections of 2-4 confocal slices of germaria from 4 day old flies for the GAL4 control (*nos*, A-A’), and germline overexpression of NLS-Act-WT (*nos>NLS-Act-WT,* B-B’) and NLS-Act-G13R monomeric actin (*nos>NLS-Act-G13R,* C-C’) stained for DAPI (A-C, blue), Fibrillarin (Fib, A-C, magenta) and FLAG to visualize the NLS-Act constructs (A-C’, green in merge). Asterisks designate the anterior tip and GSC niche in each germarium. Closed arrowheads indicate general staining patterns—structured or hazy; open arrowheads point to nuclear FLAG-actin staining within germline nuclei (B-C’). Scale bar = 20 μm. (D-F) Maximum projections of 4-6 confocal slices of whole mount ovary tile scans from 4 day old flies for the GAL4 control (*nos*, D-D’), and germline overexpression of NLS-Act-WT (E-E’) and NLS-Act-G13R (F-F’) stained for DAPI (magenta) and FLAG (green in merge). Asterisks indicate the anterior tip of each ovariole. Scale bar = 500 μm. Compared to GAL4 control germaria (A-A’), there is strong expression of structured FLAG staining in NLS-Act-WT overexpression (B-B’, arrowheads), supporting the ability for it to polymerize. NLS-Act-G13R exhibits a uniform hazy localization throughout the nuclei, consistent with this mutation preventing actin polymerization (C-C’, arrowheads). Examination of whole ovaries reveals that GAL4 control ovaries contain the expected 12-20 ovarioles and the ovarioles have the full range of stages from early to late oogenesis (D-D’); comparably, NLS-Act-WT ovaries contain the expected number of ovarioles and do not have obvious changes to the gross morphology of the ovary (E-E’). However, overexpression of NLS-Act-G13R results in ovaries that are much smaller and have less ovarioles (F-F’).

Using these two constructs, we increased wild-type or monomeric nuclear actin in the early germline and analyzed whole mount ovaries for morphological changes. A control ovary contains roughly 15-20 ovarioles, and ovarioles are composed of a range of stages (Fig. 4D-D’). Overexpressing NLS-Act-WT does not dramatically alter ovary size or composition of stages (Fig. 4E-E’), although there are subtle changes to nucleolar morphology in later stages (Talbot *et al*. 2023) and some cell death is observed at the mid-oogenesis checkpoint. Conversely, increasing monomeric nuclear actin (NLS-Act-G13R) results in ovaries that are much smaller, contain dying follicles, and have an altered stage composition, including a variable loss of early and mid-stages of oogenesis (Fig. 4F-F’). Upon closer examination, the reduced size of these ovaries appeared to be due to emptying germaria rather than a decreased number of ovarioles.

Emptying germaria result from a failure to maintain the GSC pool. During asymmetric division, if GSCs are not retained in the niche, the germarium will progressively empty as germline cysts develop and follicles form; this defect leaves behind the surrounding somatic cells and results in what is termed an ‘empty germarium’. To determine if nuclear actin contributes to GSC maintenance, we overexpressed the two forms of nuclear actin and quantified the presence of emptying/empty germarium over the course of two weeks. In control ovaries, GSCs are maintained and germaria contain GSCs, cystoblasts, and cysts (Vasa) surrounded by somatic cells (Hts) (Fig. 5A-A”). We did not observe any germline emptying over the two-week period, as expected. Like control ovaries, overexpression of polymerizable nuclear actin (NLS-Act-WT) did not result in emptying germaria (Fig. 5B-B”, F). Conversely, monomeric nuclear actin (NLS-Act-G13R) overexpression results in germaria that are at various stages of losing the germline cells (emptying) or completely empty (Fig. 5C-E”). Specifically, ovaries overexpressing monomeric nuclear actin in the germline progressively lose the germline with 9.5% empty at 2 days and 65.0% empty at 14 days of overexpression (Fig. 5F). These data suggests that nuclear actin polymerization is required for GSC maintenance.

**Figure 5.**
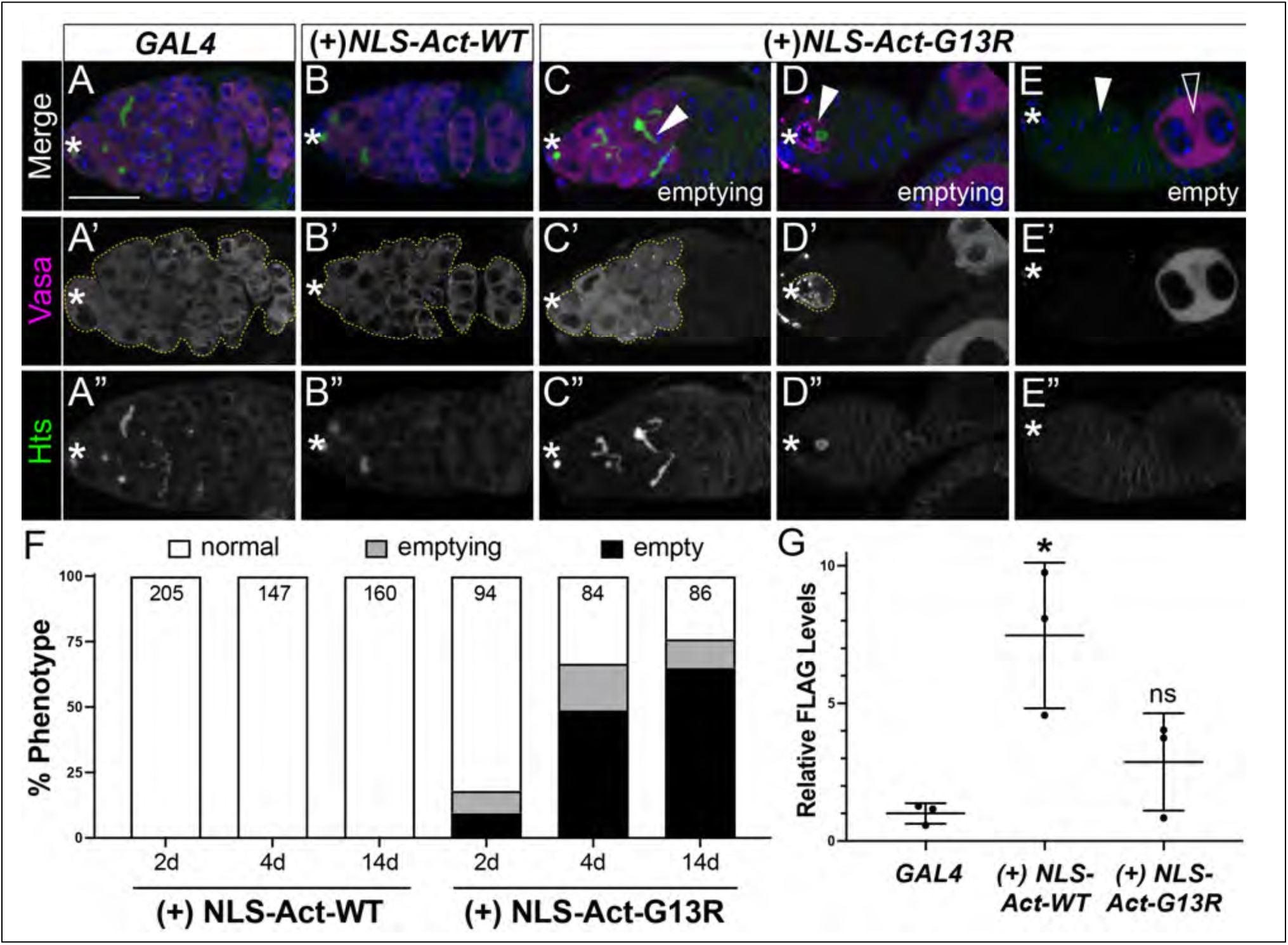
Nuclear actin polymerization is required for GSC maintenance. (A-E”) Maximum projections of 5 confocal slices of germaria of from 4 day old flies for the GAL4 control (*nos*, A-A”), and germline overexpression of NLS-Act-WT (*nos>NLS-Act-WT,* B-B”) and NLS-Act-G13R (*nos>NLS-Act-G13R,* C-E”) stained for Vasa (A-E’, magenta in merge) to marks the germline, Hts (A-E and A”-E”, green in merge) to mark the fusomes and surrounding somatic cells, and DAPI (A-E, blue) to mark the nuclei. Asterisks designate the anterior tip and GSC niche in each germarium. The germline population is outlined by yellow dashed line (A’-E’). Scale bar = 15 μm. (F) Quantification of germline loss in germaria for 2, 4, and 14 day old flies of the indicated genotypes. Germarium phenotypes were binned into three categories: normal (white), emptying (gray), and empty (black). ‘Normal’ germaria contain a range of germline cysts (GSCs through 16-cell cysts) which fill all regions of the germarium. ‘Emptying’ germaria are missing germline cysts at either the anterior or posterior end of the germarium, highlighted by regions of somatic cells absent of germline. ‘Empty’ germaria completely lack germline cells leaving behind only the surrounding somatic cells. (G) Quantification of FLAG protein levels normalized to the germline protein Vasa via Western blot for GAL4 control, NLS-Act-WT overexpression, and NLS-Act-G13R overexpression; n=3, ± SD, *, p<0.05 (p=0.0123), ns=not significant (p=0.4753). In both the GAL4 control and NLS-Act-WT germline overexpression, germaria contain GSCs and the full range of developing germline cysts (A-B’, F); the germline cysts are surrounded by somatic cells and fusomes appear normal (A”-B”). Overexpression of NLS-Act-G13R results in a range of phenotypes from partial (‘emptying’, C-D”) or complete loss (‘empty’, E-E”) of the germline. Within emptying NLS-Act-G13R overexpressing germaria there are cysts with abnormally branched fusomes (C-C”, arrowhead), and ones with a single remaining GSC, marked by a rounded fusome (D-D”, arrowhead). Empty germaria exhibit no germline cells (E-E”; white arrowhead), but may exhibit later stage follicles which sometimes have the wrong number of germline cells (open arrowhead).

It is possible the distinct phenotypes observed when polymerizable vs monomeric actin are overexpressed in the germline are due to differences in expression level of the constructs. FLAG staining of germaria expressing either NLS-actin construct are easily visible, suggesting strong expression (Fig. 4). However, the differences in the staining patterns made direct comparisons of expression levels in NLS-Act-WT versus NLS-Act-G13R difficult. Therefore, we used western blots to compare the expression of the nuclear actin constructs (Fig. 5G, S2). Both polymerizable NLS-Act-WT and monomeric actin NLS-Act-G13R constructs are expressed at robust levels, indicating that the lack of an effect on GSC maintenance by overexpression of polymerizable nuclear actin (NLS-Act-WT) is not due to low expression.

To determine if the germline loss phenotype is due to the functions of nuclear actin in early germ cells, we used additional GAL4 drivers that induce expression at different levels and times within the germline during oogenesis. First, we used *vasa*-GAL4 which drives UAS expression throughout all stages of follicle development. Similar to *nos*-GAL4, *vasa-*GAL4 overexpression of NLS-Act-G13R, but not NLS-Act-WT results in emptying germaria (Fig. S3). Further, the broader expression of this GAL4 driver led to more severe follicle death and a build-up of cellular debris under the muscle sheath (Fig. S3), indicating that polymeric nuclear actin also has roles in later stages of oogenesis. Alternatively, overexpression of NLS-Act-G13R with *mat*-GAL4, which turns on in budded follicles, does not impact GSC maintenance but does have consequences on later stages (Fig. S3). These experiments reveal a role for nuclear actin polymerization outside the germarium; however, GSC maintenance is specifically disrupted when nuclear actin polymerization is altered in the GSCs and early germline. Collectively, our overexpression experiments suggest that specific forms of actin are important for stemness and establish that nuclear actin polymerization plays a role in maintaining the germline.

### Nuclear actin is an intrinsic regulator of nuclear architecture

Nuclear size is a tightly controlled feature of stem cells to maintain stiffness, integrity of the nuclear lamina, chromatin organization, and an undifferentiated gene expression profile (Duan *et al*. 2020). We find germline cells swell beyond expected cellular and nuclear sizes when monomeric nuclear actin is overexpressed (Fig. 5D). These dramatic changes in nuclear regulation suggest that important intrinsic mechanisms maintaining cellular:nuclear size ratios are disrupted and that nuclear structures would have to re-organize to compensate for these changes. These observations led us to explore what internal nuclear component of GSCs are be regulated by nuclear actin.

We first assessed the role of nuclear actin in regulating the nuclear lamina, which is critical for maintaining the shape and boundary of the nucleus (De Leeuw *et al*. 2018). Early germline cells contain a relatively smooth, intact nuclear lamina (Lamin Dm0, Fig. 6A-A”, D white arrowheads; Fig. S4). Overexpression of NLS-Act-WT does not significantly change nuclear lamina thickness or shape, though Lamin intensity did increase compared to GAL4 control germaria (Fig. 6B-B’, E, white arrowheads; Fig. S4). When monomeric nuclear actin is overexpressed, GSCs and germline cysts exhibit significantly thickened and distorted nuclear lamina (Fig. 6C-C’, F, white arrowheads; Fig. S4). When we assessed NLS-Act-G13R germaria by the amount of germline present, we observe that Lamin intensity increases prior to losing germline cells (Fig. S4). In fact, the level of Lamin present and distortion of the lamina structure seem to negatively correlate with how much germline is present (Fig. S4). The nuclear lamina thickening is particularly striking in almost completely empty germaria and in some cases the nuclear lamina becomes extremely convoluted and breaks down (Fig. 6C-C’, F; Fig. S4). These changes to the nuclear lamina suggest altering nuclear actin, particularly nuclear actin polymerization, impacts nuclear lamina structure and can compromise nuclear integrity.

**Figure 6.**
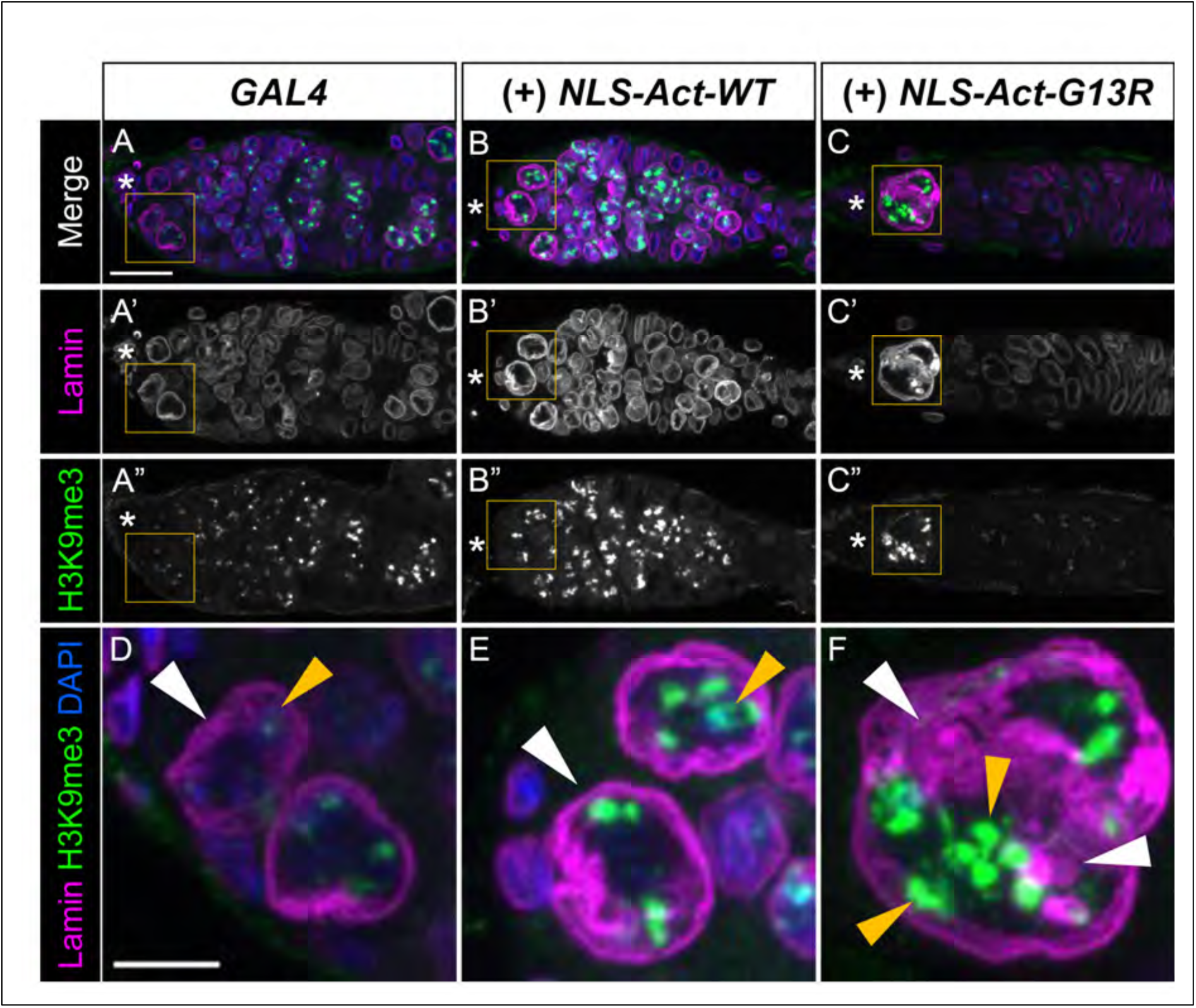
Nuclear actin polymerization is required for maintaining nuclear lamina structure and chromatin organization in the Drosophila germarium. (A-F) Maximum projections of 2-5 confocal slices of germaria from 4 day old flies for the GAL4 control (*nos*, A-A”), and germline overexpression of NLS-Act-WT (*nos>NLS-Act-WT*, B-B”) and NLS-Act-G13R (*nos>NLS-Act-G13R*, C-C”) ovaries stained for DAPI (A-C and D-F, blue in merge) to mark the nucleus, Lamin (Lamin Dm0, A-C’ and D-F, magenta in merge) to mark the nuclear lamina, and H3K9me3 (A-C, A”-C” and D-F, green in merge) to mark heterochromatin. Asterisks designate the anterior tip and GSC niche in each germarium. Panels D-F are higher magnification cropped images of yellow boxed regions of germaria in A-C. White arrowheads indicate nuclear lamina structure and yellow arrowheads indicate examples of H3K9me3 puncta. Scale bar = 15 μm in A-C”; 5 μm in D-F. Control germaria have a relatively smooth nuclear lamina (A-A’, D). Overexpression of NLS-Act-WT results in more intense Lamin staining, but no obvious loss of nuclear lamina integrity (B-B’, E). However, NLS-Act-G13R overexpression causes nuclear size to increase and nuclear lamina to thicken and exhibit distorted, convoluted regions (C-C’, F, white arrowheads). Control germaria contain H3K9me3 puncta throughout germline cells (A’A”) and a few puncta at the nuclear periphery of GSCs (A”, D, yellow arrowhead). Compared to controls, NLS-Act-WT germaria contain larger and more intense H3K9me3 puncta in both the GSCs (B-B”, E, yellow arrowhead) and the germline cells (B-B”). NLS-Act-G13R overexpression results in more, larger, and brighter heterochromatin puncta in the remaining germline cells (C, C”, F) and these puncta are found throughout the nucleoplasm (C, C”, F, yellow arrowheads) instead of at the nuclear periphery as seen in in controls (D, yellow arrowhead).

In addition to structural support, the nuclear lamina anchors chromatin to the nuclear periphery (DE LEEUW *et al*. 2018). Based on the loss of nuclear lamina integrity in germaria overexpressing NLS-Act-G13R, we hypothesized chromatin organization would be severely impacted in response to impaired nuclear actin polymerization. To visualize changes in chromatin organization, we stained germaria with an antibody to the heterochromatin mark H3K9me3. Throughout the germarium, germline cells contain distinct, small puncta of heterochromatin anchored to the nuclear envelope (Fig. 6A, A”). Wild-type GSCs contain a few puncta of heterochromatin at the nuclear periphery (Fig. 6D, yellow arrowhead). Overexpressing NLS-Act-WT moderately increases the intensity and size of H3K9me3 puncta throughout the germline (Fig. 6B, B”, and E, yellow arrowhead). However, NLS-Act-G13R overexpression increases the intensity of H3K9me3 staining beyond GAL4 controls or NLS-Act-WT overexpression and there are both more and larger puncta of heterochromatin in germline cells, including the GSCs (Fig. 6C, C”, F). These heterochromatin puncta are also distributed throughout the germline nuclei instead of being restricted to the nuclear periphery (Fig. 6F, yellow arrowheads). Similar to the changes in Lamin staining, H3K9me3 puncta become more intense and larger as NLS-Act-G13R expressing germaria increasingly lose germline cells (Fig. S4). Therefore, disrupting nuclear actin polymerization results in significant changes to the chromatin landscape of GSCs and germline cysts, which may alter the unique gene expression profiles required at each stage.

As C4 nuclear actin is also enriched in the nucleolus in the early germ cells (Wineland *et al*. 2018), we assessed nucleoli when nuclear actin is manipulated. The nucleolus is a phase-separated, membraneless organelle that forms a single rounded structure in germline cells of the germarium (Pederson 2011; Duan *et al*. 2020). Nucleolar liquid-liquid phase separation is driven by interactions of ribosomal RNA (rRNA) and proteins present in biocondensates (Lafontaine *et al*. 2021). This organizing principle means changes in nucleolar function alter nucleolar morphology; therefore, nucleolar morphology can be used as a proxy for function. To determine whether preventing actin polymerization perturbs nucleolar function and thereby, morphology, we overexpressed NLS-Act-WT and NLS-Act-G13R in the early germline and stained germaria for the nucleolar component Fibrillarin. Fibrillarin staining allows us to analyze key nucleolar morphological features including size, roundness, and the presence of fragments and/or deformations. In control germaria, each GSC and germline cell of developing cysts contain a single round nucleolus and there is an expected nucleolar size ratio of larger to smaller in GSCs through 16-cell cysts (Fig. 7A-A”, D). Like control germaria, overexpression of NLS-Act-WT results in the expected nucleolar to nuclear size ratios across the germarium and normal nucleolar morphology in GSCs (Fig. 7B-B”, E). However, in NLS-Act-G13R overexpressing germaria, both the nuclei and nucleoli of GSCs and other germline cells are swollen and larger than control nucleoli (Fig. 7C-C”, F). We also observe instances where nuclei and nucleoli of germline cells in region 2 no longer obey the expected size ratios and have comparable size ratios to those observed in GSCs (Fig. 7C-C”, arrowheads). It is atypical for single germline cells to be found in region 2, so it is unclear what the differentiation state of these cells are and what events caused their abnormal development. NLS-Act-G13R germaria also contain nucleoli which deviate from a uniform round shape and are sometimes fragmented (Fig. S5, arrowheads). Like changes to the nuclear lamina and H3K9me3 puncta, nucleolar morphology is altered before germline loss begins and becomes more severe as germline cells are lost (Fig. S5). These data indicate nuclear actin polymerization prevents nucleolar hypertrophy and is important for maintaining nuclear and nucleolar size ratios as well as normal nucleolar morphology.

**Figure 7.**
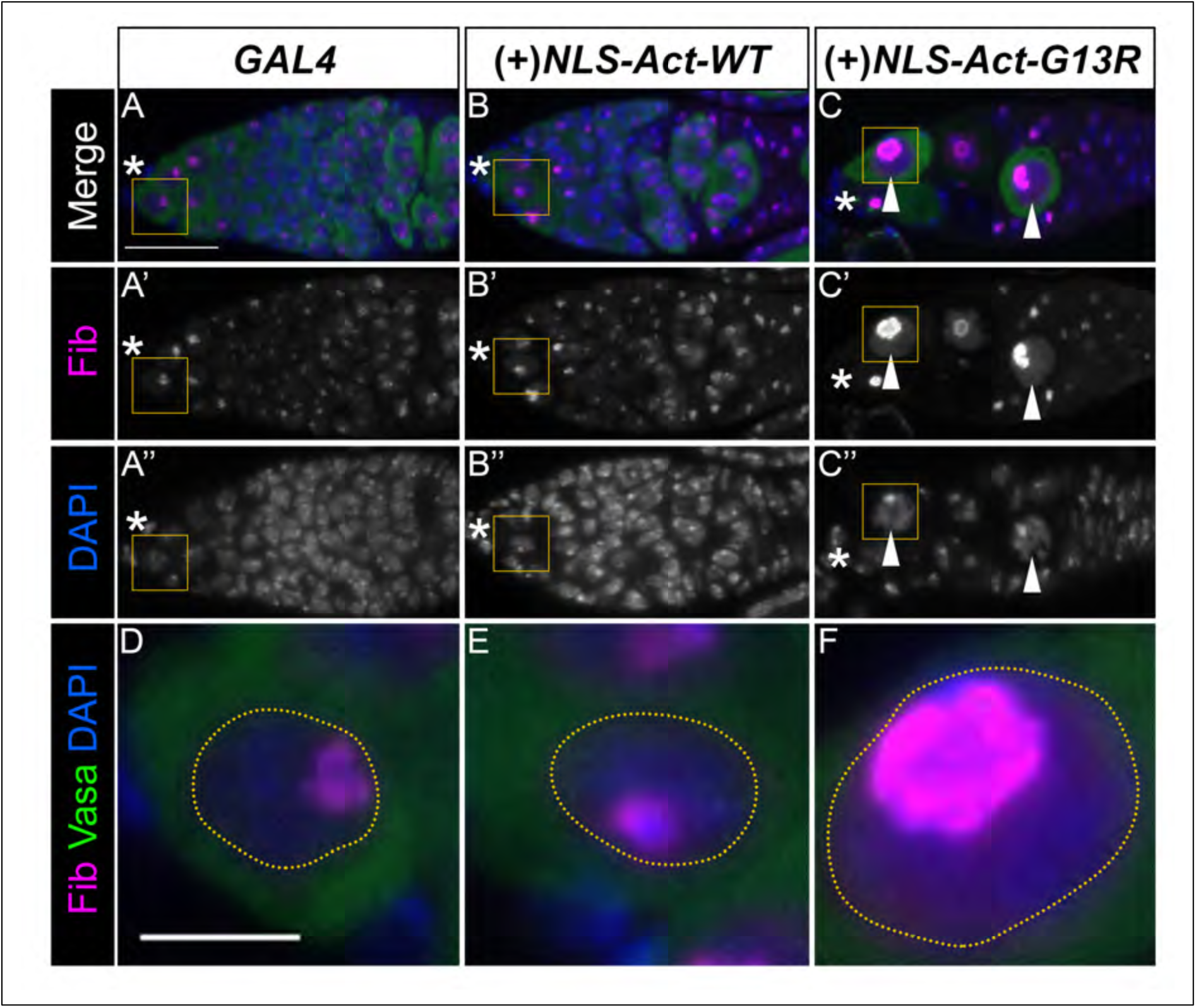
Nuclear actin polymerization regulates nucleolar morphology in the early germline. (A-F) Maximum projections of 2-5 confocal slices of germaria from 4 day old flies for the GAL4 control (*nos*, A-A”), and germline overexpression of NLS-Act-WT (*nos>NLS-Act-WT*, B-B”) and NLS-Act-G13R (*nos>NLS-Act-G13R*, C-C”) stained for DAPI (A-F, A”-C”, blue in merge) to mark the nucleus, Fibrillarin (Fib, A-F, A’-C’, magenta in merge) to mark the nucleolus, and Vasa (A-F, green in merge) to mark germline cytoplasm. Asterisks designate the anterior tip and GSC niche in each germarium. Panels D-F are higher magnification cropped images of yellow boxed regions of germaria in A-C. Scale bar = 15 μm in A-C”; 5 μm in D-F. Control GAL4 germaria have specific nucleolar (Fib, A-A’, magenta) and nuclear (DAPI, A”-B”, blue) sizes, where GSCs have larger nucleoli and nuclei and these size ratios decrease as cells become more differentiated (2-, 4-, 8-, 16-cell cysts) until becoming larger again in regions 2b-3. Whereas NLS-Act-WT germaria show the expected nuclear and nucleolar sizes (B-B”), NLS-Act-G13R germaria have germline cells with nuclear and nucleolar hypertrophy (arrows, C-C”). In NLS-Act-G13R germaria that are emptying, individual germline cells improperly located in region 2b are similarly sized, displaying a loss of the expected nuclear size trends present in control germaria (A-A’”). Higher magnification images of single GSC nuclei show similar nuclear and nucleolar sizes in GAL controls (yellow dashed line, D) and NLS-Act-WT (yellow dashed line, E), but distinctly larger, swollen nuclei and nucleoli (yellow dashed line, F) after NLS-Act-G13R overexpression.

Given that perturbing the balance of polymeric and monomeric nuclear actin results in striking alterations in the nuclear lamina, heterochromatin, and nucleoli, and that these three nuclear architecture components likely impact each other (Duan *et al*. 2020), we next sought to determine which defects arise first. Specifically, we assessed these three nuclear architecture components after 24 hours of NLS-Act-G13R overexpression. However, we could not distinguish a nuclear structure that is preferentially affected or a distinctive order in which nuclear architecture is impacted (data not shown). Rather, GSC nuclei either appeared normal or all three components showed subtle or more pronounced changes in NLS-Act-G13R overexpressing germaria. Further studies will be required to dissect the complex interactions between the nuclear architecture components and to determine whether nuclear actin has direct or indirect roles in maintaining each of these nuclear components. However, our data indicate that nuclear actin polymerization plays an essential role in the intrinsic regulation of several key nuclear architecture components within the GSCs and the developing germline.

### Restoring the balance of polymeric and monomeric nuclear actin pools rescues germline loss

Our data lead to the model that a specific distribution of polymeric and monomeric actin is required in GSCs for their maintenance, and changes in this distribution are required for proper germ cell differentiation. Specifically, our data supports that nuclear actin polymerization is required for GSC functions, regulates nuclear architecture, and maintains stemness. If this is true, then supplying polymerizable nuclear actin in the germline of flies overexpressing NLS-Act-G13R should suppress germline loss.

To test this idea, we simultaneously overexpressed polymerizable NLS-Act-WT and NLS-Act-G13R (*nos>NLS-Act-WT; NLS-Act-G13R*). We first verified that both constructs are expressed. We observe a more structured actin staining in polymerizable NLS-Act-WT germaria (Fig. 8B-B’; Fig. S5) and a hazy, unstructured pattern in NLS-Act-G13R germaria where germline is still present (Fig. 8C-C’; Fig. S5). In our co-overexpression system, we see both structured FLAG staining at the cell cortex and also a haze of FLAG-actin staining throughout the nucleus (Fig. 8E-E’; Fig. S5), indicating both monomeric and polymeric nuclear actin are present. We next assessed whether re-introducing polymerizable nuclear actin via our co-overexpression system could rescue the emptying/empty germaria observed in NLS-Act-G13R ovaries. After four days of overexpression, neither GAL4 controls nor NLS-Act-WT overexpression alone show any germline loss (Fig. 8A-A”, G). Similar to the data in Figure 5, overexpression of only NLS-Act-G13R causes 47.7% of germaria to completely lose the germline (Fig. 8C-D”). Strikingly, co-overexpression of NLS-Act-G13R and polymerizable NLS-Act-WT results in a dramatic rescue of emptying germaria, with only 2.1% completely empty and 12.1% emptying germaria (Fig. 8E-F”, G). Further, re-introducing polymerizable nuclear actin in the NLS-Act-G13R background restores a more normal GSC and germline cyst cell and nuclear size ratios (DAPI, Fig. 8A’’’-F’’’; Fig. S6). However, we do observe some NLS-Act-WT; NLS-Act-G13R germaria with deformed nucleoli, despite expected nuclear:nucleolar ratios (Fig. S6). Overall, restoring the ability of nuclear actin to polymerize results in a robust improvement in germline maintenance. Our findings establish the importance of polymeric nuclear actin in GSC maintenance processes and in regulating the internal architectures of GSC nuclei.

**Figure 8.**
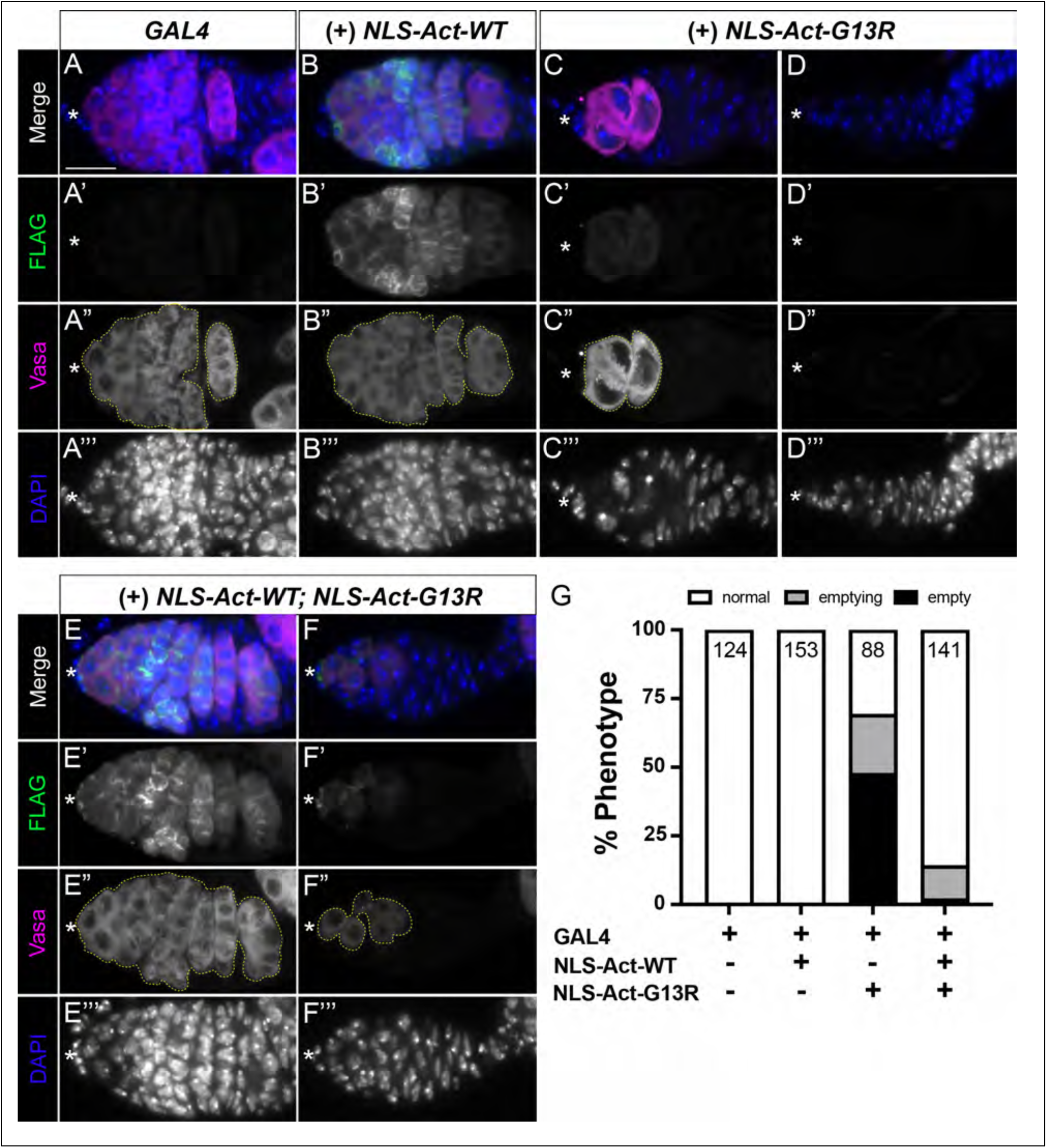
Restoring nuclear actin polymerization rescues germline loss in NLS-Act-G13R overexpression. (A-F’’’) Maximum projections of 2-5 confocal slices of germaria from 4 day old flies for the GAL4 control (*nos*, A-A’’’), and germline overexpression of polymerizable NLS-Act-WT (*nos>NLS-Act-WT*, B-B’’’), monomeric NLS-Act-G13R (*nos>NLS-Act-G13R*, C-D’’’), and of both polymerizable NLS-Act-WT and monomeric NLS-Act-G13R (*nos>NLS-Act-WT; NLS-Act-G13R*, E-F’’’) stained for FLAG (A-F’, green in merge), Vasa to label the germline (A-F, A”-F”, magenta in merge), and DAPI to label the nucleus (A-F, A’’’-F’’’, blue in merge). Asterisks designate the anterior tip and GSC niche in each germarium. Yellow dotted lines outline germline content in each germarium (A”-F”). Scale bar = 15 μm. (G) Quantification of germline loss in germaria from 4 day old flies. Germarium phenotypes were binned into three categories: normal (white), emptying (gray), and empty (black). ‘Normal’ germaria contain a range of germline cysts (GSC through 16 cell cysts) which fill all regions of the germarium. ‘Emptying’ germaria are missing germline cysts at either the anterior or posterior end of the germarium, highlighted by regions of somatic cells absent of germline. ‘Empty’ germaria are completely absent of germline cells leaving behind only the surrounding somatic cells. FLAG staining is absent in GAL4 controls as expected (A-A’, arrowhead). In NLS-Act-WT overexpression, FLAG staining is structured indicating the ability to polymerize (B-B’, arrowhead), whereas NLS-Act-G13R FLAG staining is hazy (C-C’, arrowhead) and diffuse consistent with the G13R mutation blocking polymerization. In the latter case, as germline cells are lost, FLAG staining decreases (C’, arrowheads; also see Supplemental Figure 5). Co-overexpression of NLS-Act-WT and NLS-Act-G13R results in structured FLAG staining (E-F’, arrowheads), suggesting nuclear actin polymerization is restored. In both control GAL4 and NLS-Act-WT overexpression, the germaria contain GSCs and the full range of developing germline cysts (A-B’’’, G). Overexpression of NLS-Act-G13R results in emptying germaria (C, C”, G), often with germline cells having larger than expected nuclei (C’’’, yellow outline), or empty germaria that are comprised of only somatic cells (D-D”, G). Co-overexpression of polymerizable nuclear actin and monomeric nuclear actin results in primarily ‘normal’ germaria (E-E”), dramatically reducing the presence of emptying or empty germaria (F-F”) in comparison to overexpression of monomeric nuclear actin (NLS-Act-G13R) alone (G).

## DISCUSSION

Using *Drosophila* female GSCs as a model, we provide the first evidence that the balance of monomeric and polymeric nuclear actin is critical for controlling stemness versus differentiation in an *in vivo*, physiological context. C4 nuclear actin localization and form change during *Drosophila* germline cell differentiation. The nucleoplasmic C4 nuclear actin present in the GSCs and early germ cells is lost as differentiation proceeds. Whereas monomeric C4 nuclear actin labels the nucleoli from the GSCs through the 8-cell cysts. By overexpressing NLS-actin constructs – either monomeric or wild-type – we find that polymeric nuclear actin is required to maintain GSC fate and proper germline development. Overexpression of monomeric nuclear actin results in GSC, and thereby, germline, loss. Potentially contributing to the GSC loss, we find that overexpression of monomeric nuclear actin results in nuclear and nucleolar hypertrophy, increased heterochromatin, and alteration of the nuclear lamina. Supporting that these phenotypes are due to the polymerization state of nuclear actin, co-overexpression of both monomeric and wild-type nuclear actin suppresses all the defects. Together, these data lead to the model that tight regulation of nuclear actin polymerization is required for the nuclear organization and related functions necessary for maintaining GSC fate and subsequent germline differentiation.

### Connecting C4 nuclear actin dynamics to germ cell differentiation

We find the nucleoplasmic, polymeric C4 nuclear actin is present in the GSCs, cystoblasts and 2-cell cysts, but is absent from the 4-cell cysts onward. This pattern led us to assess the roles of nuclear actin in early germ cell fate, but also suggests the 2-cell to 4-cell cyst transition represents a key step in differentiation. Supporting this idea, only the first two germ cells (those comprising the 2-cell cyst) can become the oocyte, a true totipotent cell. At the 4-cell cyst stage mitosis begins – the synaptonemal complex forms and oocyte markers become enriched the two pro-oocytes (Hinnant *et al*. 2020). These data are consistent with polymeric C4 nuclear actin marking the less differentiated germ cells. Further supporting that polymeric nuclear actin marks undifferentiated cells, C4 polymeric nuclear actin is highly enriched in the differentiated oocyte starting from Region 3 of the germarium (Wineland *et al*. 2018). Additionally, the fusome structure changes from linear in the 2-cell cysts to branched in 4-cell cysts. Differences in the fusome across the cells within a cyst contribute to oocyte fate and the differentiation of the other germ cells into nurse cells (Hinnant *et al*. 2020). Notably, the fusome is a cytoskeletal structure and is enriched for F-actin. As changes in cytoplasmic actin can regulate nuclear actin levels and structure (Kelpsch and Tootle 2018; Ulferts *et al*. 2021), it is tempting to speculate that the fusome changes contribute to the changes in nuclear actin form, and these changes contribute to the differentiation of the germline.

Other evidence supporting similarities in cell fate from the GSCs to the 2-cell cysts come from single cell RNA sequencing (scRNAseq) studies. Two studies clustered the GSCs, cystoblast, and 2-cell cysts together (Rust *et al*. 2020; Slaidina *et al*. 2021). This clustering indicates a similarity in gene expression but could be an artifact due to the scarcity of these cells. However, striking expression changes are apparent between the 2-cell and 4-cell cysts (Rust *et al*. 2020), supporting that the differentiation state of these cells may be different.

Additionally, the early germ cells are wrapped by somatic cells termed escort cells (also referred to as inner germarial sheath cells), and these cells play critical roles in regulating germline differentiation (Hinnant *et al*. 2020; Giedt and Tootle 2023). scRNAseq data supports there are multiple classes of escort cells, ranging from two to three groups (Rust *et al*. 2020; Shi *et al*. 2021; Slaidina *et al*. 2021; Tu *et al*. 2021). In all the studies the anterior escort cells are distinct from the posterior ones, but two studies indicate a third, central group of escort cells (Rust *et al*. 2020; Slaidina *et al*. 2021). Intriguingly, the anterior to central escort cell distinction occurs around the 2- to 4-cell transition, raising the possibility that differences in signaling from the distinct populations of escort cells contribute to the change in nuclear actin forms and the differentiation of the cysts. Together, these data support that the 2-cell to 4-cell transition is a key step in germline differentiation, and our work implicates changes in nuclear actin structure as a key contributor.

### Nuclear actin – a conserved regulator of cell fate

Multiple studies support that tight control of nuclear actin is essential for controlling cell fate and differentiation. For example, mouse embryonic fibroblasts (MEFs) from β-actin knockout animals have increased repressive histone modifications that block neuronal reprogramming; expression of nuclear targeted β-actin restores differentiation (Xie *et al*. 2018a; Xie *et al*. 2018b). Follow-up studies reveal β-actin modulates the activities of multiple chromatin remodeling complexes, to impact both local chromatin accessibility and 3D genome architecture to regulate gene expression necessary for development and differentiation (Mahmood *et al*. 2021). These studies reveal nuclear actin plays critical roles in determining the gene expression and therefore, the differentiation state of cells. However, these studies do not address what form of nuclear actin is involved (monomers or polymers), leaving it unclear how nuclear actin structure impacts cell fate.

We find that polymeric nuclear actin is required for *Drosophila* GSC maintenance, as overexpression of monomeric nuclear actin results in GSC loss and subsequent, germline loss. This is likely a conserved function of polymeric nuclear actin, at least in certain cell types. Indeed, reprogramming of somatic nuclei transplanted into Xenopus oocytes requires nuclear actin polymerization (Miyamoto *et al*. 2011a). Blocking nuclear actin polymerization, by actin antibody injection or inhibitor treatment, prevents *Oct4*-dependent reprogramming, and expression of polymerization competent, but not monomeric actin mutants (including G13R), promotes *Oct4* expression. Further, studies on cultured mesenchymal stem cells reveal their differentiation into osteoblasts or adipocytes is regulated by the level and structure of nuclear actin (Sankaran *et al*. 2019a; Rubin *et al*. 2022). In particular, a stiff environment induces cytoskeletal remodeling, which promotes the nuclear accumulation of actin, driving polymerization by mDia2, a nuclear localized actin elongation factor (Sen *et al*. 2011; Sen *et al*. 2015a; Sen *et al*. 2017; Sen *et al*. 2020). This accumulation of nuclear polymeric actin alters the nucleoskeleton and chromatin organization, increasing nuclear stiffness, and ultimately, resulting in an osteogenic fate. Conversely, inhibiting Arp2/3, a branched actin nucleating complex, promotes adipogenesis. Recent work reveals specific changes in actin filament organization result in changes to chromatin accessibility to drive different the differentiation states (Sen *et al*. 2024). These studies indicate that actin binding proteins localize to the nucleus and promote nuclear actin polymerization, and this polymeric actin regulates nuclear structure and chromatin organization to control cell fate. It will be important to determine the mechanisms promoting nuclear actin polymerization in the *Drosophila* GSCs, including the actin binding proteins and the signaling pathways regulating them.

### Nuclear actin and nuclear architecture

Like the studies above, we find that perturbing the structure of nuclear actin within the *Drosophila* GSCs results in striking changes in nuclear structure and chromatin state; additionally, we find dramatic alterations in nucleolar size and shape. There are a number of ways in which altering the polymerization state of nuclear actin could result in these defects.

Nuclear structure is regulated by both the nucleoskeleton and chromatin (Stephens *et al*. 2019). The nucleoskeleton is comprised of the nuclear lamina – a network of lamin intermediate filaments and lamin-associated proteins including Emerin – and actin (Adam 2017; Gunasekaran *et al*. 2022). *In vitro* studies find that lamins can bind and bundle F-actin (Sasseville and Langelier 1998; Simon *et al*. 2010). Supporting this interaction may be physiologically relevant, studies using *Drosophila* cultured cells found that knockdown of the A-type lamin increases nuclear actin polymerization (Dopie *et al*. 2015). Emerin is also implicated in regulating actin. Specifically, Emerin acts as a pointed end capping protein and increases actin polymerization rates *in vitro* (Holaska *et al*. 2004), and it is speculated that this regulation of nuclear F-actin mediates the localization of chromatin remodeling complexes to the nuclear periphery (Holaska and Wilson 2007). The nuclear lamina binds to chromatin, anchoring closed and heterochromatic regions to the nuclear periphery. Recent work reveals that chromatin state also impacts the structure and stiffness of the nucleus. Indeed, decreasing heterochromatin results in nuclear deformations and softening, whereas increased heterochromatin stiffens nuclei (Stephens *et al*. 2017; Stephens *et al*. 2018). As monomeric nuclear actin functions in chromatin remodeling complexes (Klages-MUNDT *et al*. 2018) and inhibits histone deacetylase activity (Serebryannyy *et al*. 2016), regulating nuclear actin polymerization is critical for controlling chromatin state. Thus, nuclear actin can contribute to nuclear structure by regulating chromatin and by interacting with the nuclear lamina components, which in turn may control nuclear actin polymerization.

Nuclear stiffness changes with differentiation, with stem cells having softer nuclei than their differentiating daughters (Swift and Discher 2014). Indeed, the *Drosophila* GSCs have a different nucleoskeletal composition and chromatin organization than the differentiating cysts (Duan *et al*. 2020). Perturbing the nucleoskeleton of the GSC induces a quality control checkpoint and results in GSC loss (Barton *et al*. 2013; Barton *et al*. 2016; Barton *et al*. 2018). We speculate that the nucleoskeleton of the GSCs – the lamins and Emerin – regulate nuclear actin polymerization and polymeric nuclear actin limits chromatin remodeling by reducing monomer concentration, ensuring that the differentiation program remains silent. Given the severe changes in GSC nuclear structure when monomeric nuclear actin is increased, it seems likely that polymeric nuclear actin also modulates nucleoskeletal structure.

We also find that that altering GSC nuclear actin results in severe changes in nucleolar shape and size. Monomeric nuclear actin localizes to the nucleoli of the early germ cells. The nucleolus is the site of rRNA transcription, and forms by phase separation, meaning that alterations in function cause changes in structure (Pederson 2011; Lafontaine *et al*. 2021). As actin is a co-factor for RNA polymerases (RNAPs) (Kelpsch and Tootle 2018; Sokolova *et al*. 2018; Green *et al*. 2021), including RNAPI in the nucleolus (Percipalle 2013), we speculate nuclear actin plays a critical role in regulating nucleolar transcription in the early germ cells; however, the roles of polymeric versus monomeric nuclear actin remain unclear. Monomeric nuclear actin may also modulate rRNA processing and transport out of the nucleus (Percipalle *et al*. 2002; Obrdlik *et al*. 2008; Percipalle 2013). Nucleolar functions may also be impacted by nuclear actin-dependent alterations in chromatin, as heterochromatin is associated with the nucleolus and regulates both its transcription and phase separation (Lafontaine *et al*. 2021). Further, in later stages of *Drosophila* oogenesis, increasing polymerize-competent nuclear actin increases nucleolar transcription which drives changes in nucleolar structure, and increases protein translation (Talbot *et al*. 2023). These data reveal nuclear actin has multiple functions that impinge on regulating the nucleolus.

Controlling nucleolar activity is critical for stem cell maintenance and differentiation (Buszczak *et al*. 2014). The *Drosophila* GSCs have a large nucleolus, and nucleolar size decreases with differentiation (Duan *et al*. 2020). This size difference positively correlates with rRNA transcription levels and decreasing rRNA transcription or processing results in GSC loss (Neumuller *et al*. 2013; Zhang *et al*. 2014; Sanchez *et al*. 2016). Despite the high level of rRNA produced in the GSCs, ribosome biogenesis and protein synthesis are low (Zhang *et al*. 2014; Sanchez *et al*. 2016); a similar dichotomy is observed in embryonic stem cells versus their differentiated embryoid bodies (Buszczak *et al*. 2014). In *Drosophila*, knockdown of ribosomal assembly factors results in aberrant germ cell differentiation and the formation of stem cysts (Sanchez *et al*. 2016). Given the increased size and abnormal shape of the early germ cell nucleoli when monomeric nuclear actin is increased, it is likely that misregulation of rRNA transcription and protein translation contribute to the GSC loss.

## Conclusion

Based on the above findings, in conjunction with this study, we favor the model that the balance of monomeric to polymeric nuclear actin is both sensed and controlled by multiple nuclear components; when the balance is perturbed it induces cellular stress responses that when prolonged lead to GSC loss. Indeed, misbalance of nuclear actin in the GSCs impairs nucleolar functions, and this could activate the nucleolar stress response (Pederson 2011; NUNEZ Villacis *et al*. 2018). Similarly, we postulate the nuclear actin misbalance is sensed by the nuclear lamina components, leading to changes in nuclear structure; such alterations activate the GSC quality control checkpoint (Barton *et al*. 2018). Notably, the same factors, ATR and Chk2, drive both the GSC quality control checkpoint and the nucleolar stress response (Pederson 2011; NUNEZ Villacis *et al*. 2018); whether these factors contribute to the GSC loss when polymeric nuclear actin is reduced remains to be determined. The misbalance of nuclear actin, by perturbing the nucleoskeleton, the nucleolus, and/or chromatin remodeling factors directly, also drives alterations in chromatin state. While the order of these effects remains unclear, any one of these changes alone could drive the GSC loss observed when nuclear actin forms are misbalanced. Given the conserved roles of nuclear actin across organisms, we speculate that the proper balance of monomeric to polymeric nuclear actin is a critical regulator of stemness and differentiation across tissues and organisms.

## DATA AVAILABILITY STATEMENT

All data for this study are provided in the article and supplementary files. Inquiries can be directed to the corresponding author.

## AUTHOR CONTRIBUTIONS

NMG and DET performed experiments and analyzed data. NMG performed statistical analyses and designed figures. NMG and TLT conceived the study. TLT secured funding and administered the project. NMG and TLT wrote and revised the manuscript.

## FUNDING

This project was supported by National Science Foundation MCB2017797 (TLT), National Institutes of Health (NIH) R01 GM116885 (TLT), and NIH R35 GM144057. NMG was supported by a Postdoctoral Teaching Fellowship from the Department of Anatomy & Cell Biology, Carver College of Medicine, University of Iowa. DET was supported by a Summer Fellowship from the Graduate College, University of Iowa.

## ACKNOWLEDGEMENTS

We are grateful to Michael Buszczak, Pamela Geyer, Maria Vartiainen, and Peter Vilmos for Drosophila stocks and reagents. We thank the Westside Fly Group, and the Dunnwald, Geyer, and Wallrath labs for helpful discussions, and members of the Tootle Lab for helpful discussions and careful review of the manuscript. Stocks obtained from the Bloomington Drosophila Stock Center (NIH P40OD018537) were used in this study. At the University of Iowa, Information Technology Services–Research Services provided data storage support.

## CONFLICT OF INTEREST

The authors declare that the research was conducted in the absence of any commercial or financial relationships that could be construed as a potential conflict of interest.

The authors declare no competing or financial interests.

## Supplemental Figures and Legends

**SFigure 1.**
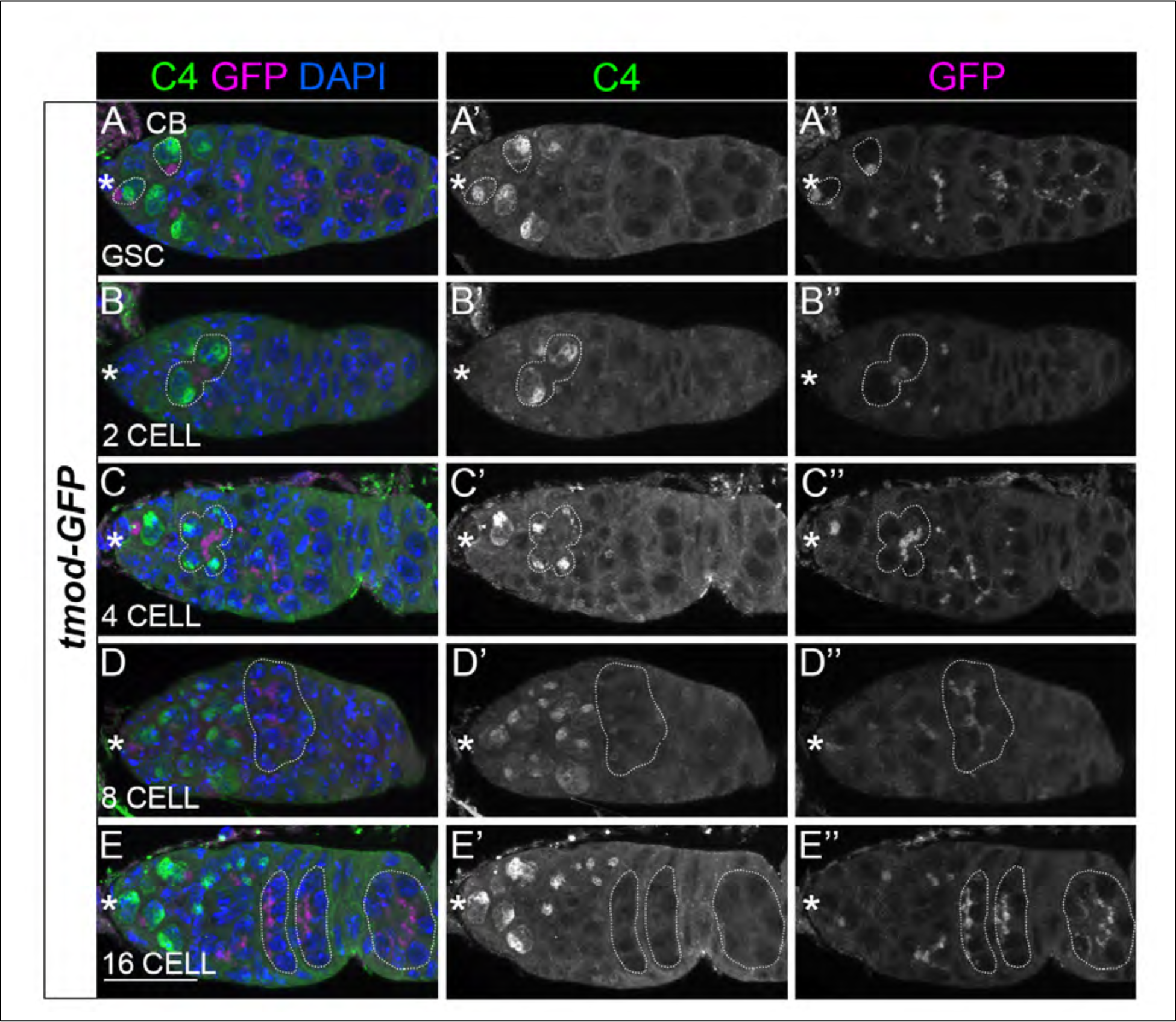
C4 nuclear actin characterization via cyst staging with tmod-GFP reporter. (A-E”) Maximum projections of 5-10 confocal slices of germaria from 4 day old flies expressing *tmod-GFP* stained for DAPI (blue, A-E), C4 actin (green, A-E’), and GFP (magenta, A-E, A”-E”). Anti-actin C4 staining is present in both the cytoplasm and in the nucleus (A-E). Tmod-GFP marks the fusomes which can be used to distinguish stages of germline development (A-E, A”-E”). GSCs and cysts are outlined by white dashed lines as indicated. Asterisks designate the anterior tip and GSC niche in each germarium. Scale bar 15=μm. Consistent with stage-specific localization shown in Figure 3, the nucleoplasmic C4 pool is present in GSCs through 2-cell cysts (A-B”) and the nucleolar pool is present in GSCs through 8-cell cysts (A-D”) and is absent in 16-cell cysts (E-E”). Note that C4 staining is absent in both lens phase 16-cell cysts and encapsulated 16-cell cysts.

**SFigure 2.**
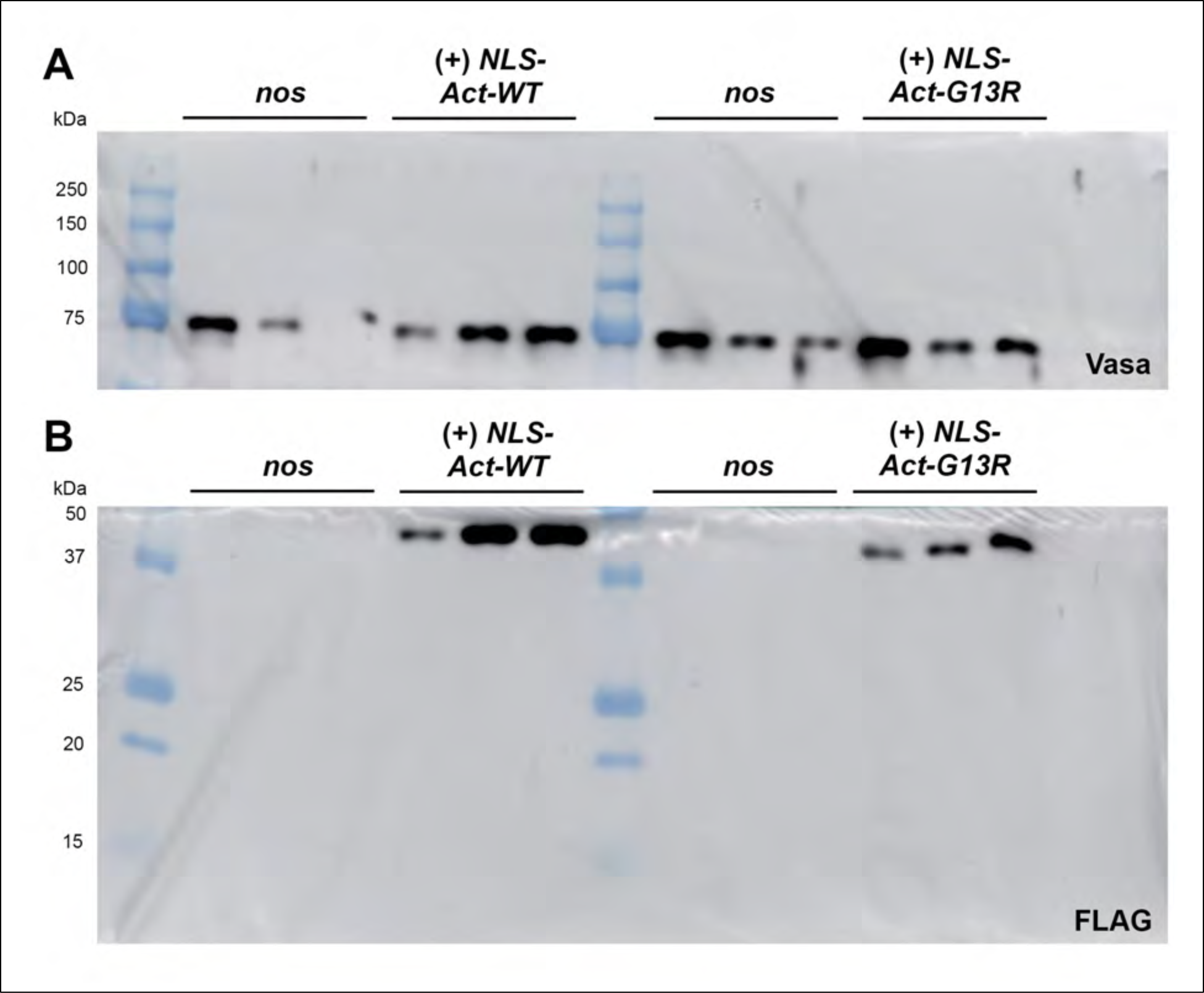
Western blot for NLS-FLAG-Act expression. (A-B) Western blots of whole ovary lysates of control (*nos>yw*), wild-type nuclear actin overexpression (*nos>NLS-Act-WT*), and monomeric nuclear actin overexpression (*nos>NLS-Act-G13R*). Blot was cut after transfer and stained with either Vasa (A, loading control) or FLAG (B, NLS-Act overexpression). Western blot of control stained for Vasa (loading control) used to normalize NLS-Act levels in quantification (Figure 5G). FLAG staining is absent in control ovaries and both NLS-Act-WT and NLS-Act-G13R are adequately expressed in ovaries (B, also see Figure 5). Molecular weight ladder (BioRad Precision Plus Protein Standard) is labeled in kDa measurements on each blot for reference.

**SFigure 3.**
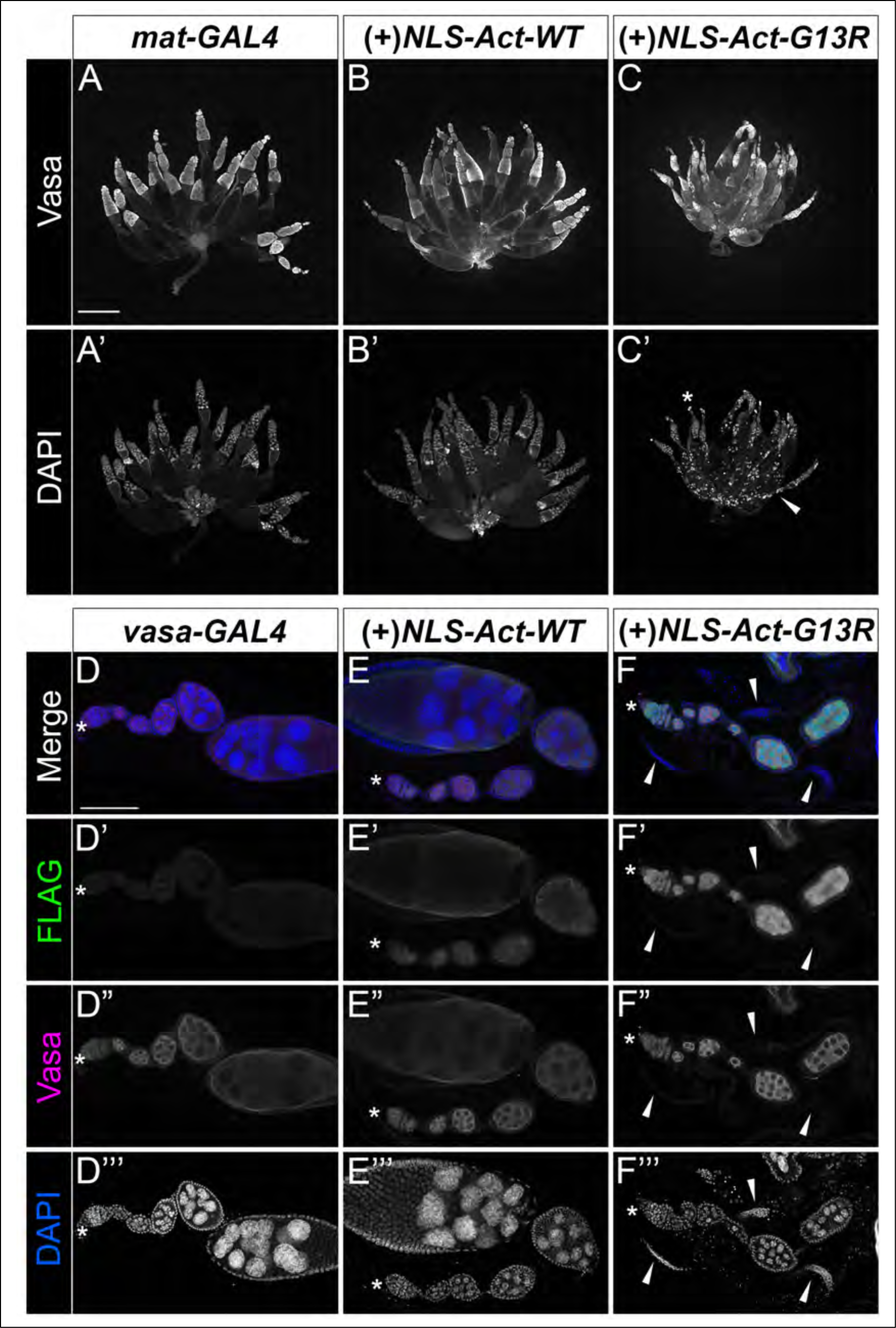
NLS-Act-G13R germline overexpression at different stages of development results in distinct phenotypes. (A-C’) Maximum projections of 4-6 confocal slices of whole mount ovary tile scans from 4 day old flies for the *mat-GAL4* control (A-A’), NLS-Act-WT overexpression (B-B’), and NLS-Act-G13R monomeric actin overexpression (C-C’) stained for Vasa (A-C) to label the germline and DAPI (A’-C’) to label the nuclei of both germline and somatic cells. Scale bar = 400 μm. (D-F’’’) Maximum projections of 2-3 confocal slices of ovarioles from 4 day old flies for the *vasa-GAL4* control (D-D’’’), NLS-Act-WT overexpression (E-E’’’), and NLS-Act-G13R overexpression (F-F’’’) stained for FLAG (green in merge), Vasa (magenta in merge), and DAPI (blue in merge). Asterisks designate the anterior tip and GSC niche in each germarium. Arrowheads indicate emptying/empty germaria. Scale bar = 100 μm. When NLS-Act-WT (B-B’) and NLS-Act-G13R (C-B’) are expressed using the *mat-*GAL4 driver (A-A’), the germline is present. However, NLS-Act-G13R overexpression results in condensed DAPI puncta in mid- to late oogenesis (C’, arrowhead), and nuclei accumulate under the muscle sheath at tips of ovarioles in NLS-Act-G13R ovaries (C’, asterisk), indicative of dying follicles. Using the *vasa-GAL4* driver, NLS-Act-WT exhibits a structured appearance (E-E’) and NLS-Act-G13R appears as a nuclear haze (F-F’; compare to similar FLAG staining with *nos-GAL4* overexpression in Fig. 4). Germline cells are lost in the germaria and ovarioles of NLS-Act-G13R overexpression using *vasa-*GAL4 (F-F’’’, arrowheads), but not in GAL4 controls (D-D’’’) or NLS-Act-WT overexpression (E-E’’’).

**SFigure 4.**
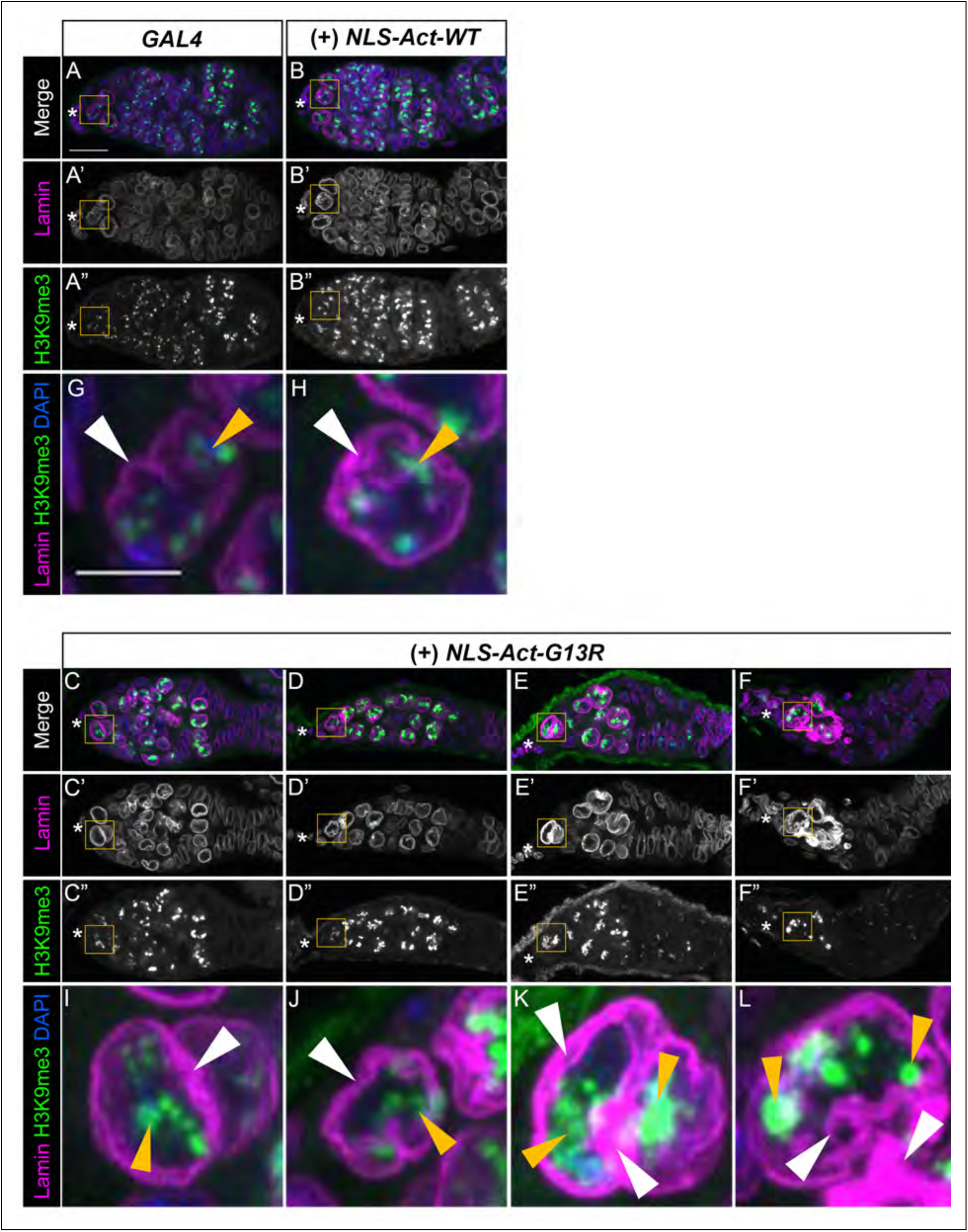
Additional examples of nuclear lamina and chromatin structure in control versus NLS-Act overexpression germaria. (A-F) Maximum projections of 2-5 confocal slices of germaria from 4 day old flies for the GAL4 control (*nos*, A-A”), and germline overexpression of NLS-Act-WT (*nos>NLS-Act-WT*, B-B”) and NLS-Act-G13R (*nos>NLS-Act-G13R*, C-F”) ovaries stained for DAPI (blue in merge) to mark the nucleus, Lamin (Lamin Dm0, magenta in merge) to mark the nuclear lamina, and H3K9me3 (green in merge) to mark heterochromatin. Asterisks designate the anterior tip and GSC niche in each germarium. The range of nuclear lamina and heterochromatin phenotypes observed in NLS-Act-G13R germaria are shown in C-F”, with emptying phenotype presented from least to most severe. (G-L) Cropped higher magnification of GSCs from yellow boxed regions of germaria in A-F. White arrowheads point out nuclear lamina and yellow arrowheads indicate examples of heterochromatin puncta. Scale bar = 15 μm in A-F”; 5 μm in G-L. Control germaria have a uniform, continuous nuclear lamina (A, A’, G). Overexpression of NLS-Act-WT leads to increased intensity of Lamin staining throughout all germline cysts (B, B’), including GSCs (H), though there are no dramatic changes to nuclear lamina structure (H, white arrowhead). Alternatively, in NLS-Act-G13R overexpressing germaria, nuclear lamina thickening and distortion correlates with increases in germline nuclei size as germaria progressively empty (left to right, C-F’’ and I-L, white arrowheads). Control germaria contain H3K9me3 puncta throughout germline cells (A, A”) and a few small, round puncta at the nuclear periphery of GSCs (A”, B, yellow arrowhead). Compared to controls, NLS-Act-WT overexpressing germaria contain larger H3K9me3 puncta in GSCs and germline cells (B, B” and H, yellow arrowhead). NLS-Act-G13R overexpression also results in more, larger, and more intense H3K9me3 puncta in germline cells (C-F” and I-L, yellow arrowheads). Even in NLS-Act-G13R germaria that have not begun to lose germline cells, H3K9me3 puncta in GSCs are larger and have a more internal localization (C-C” and I, yellow arrowhead) compared to GAL4 controls where H3K9me3 puncta are located at the nuclear periphery (A-A”, and G, yellow arrowhead). As germline cells are lost from NLS-Act-G13R germaria (D”-F” and J-L), H3K9me3 puncta grow in size and intensity, sometimes coalescing into large H3K9me3 territories occupying significant regions of GSC nuclei (K-L, yellow arrowheads).

**SFigure 5.**
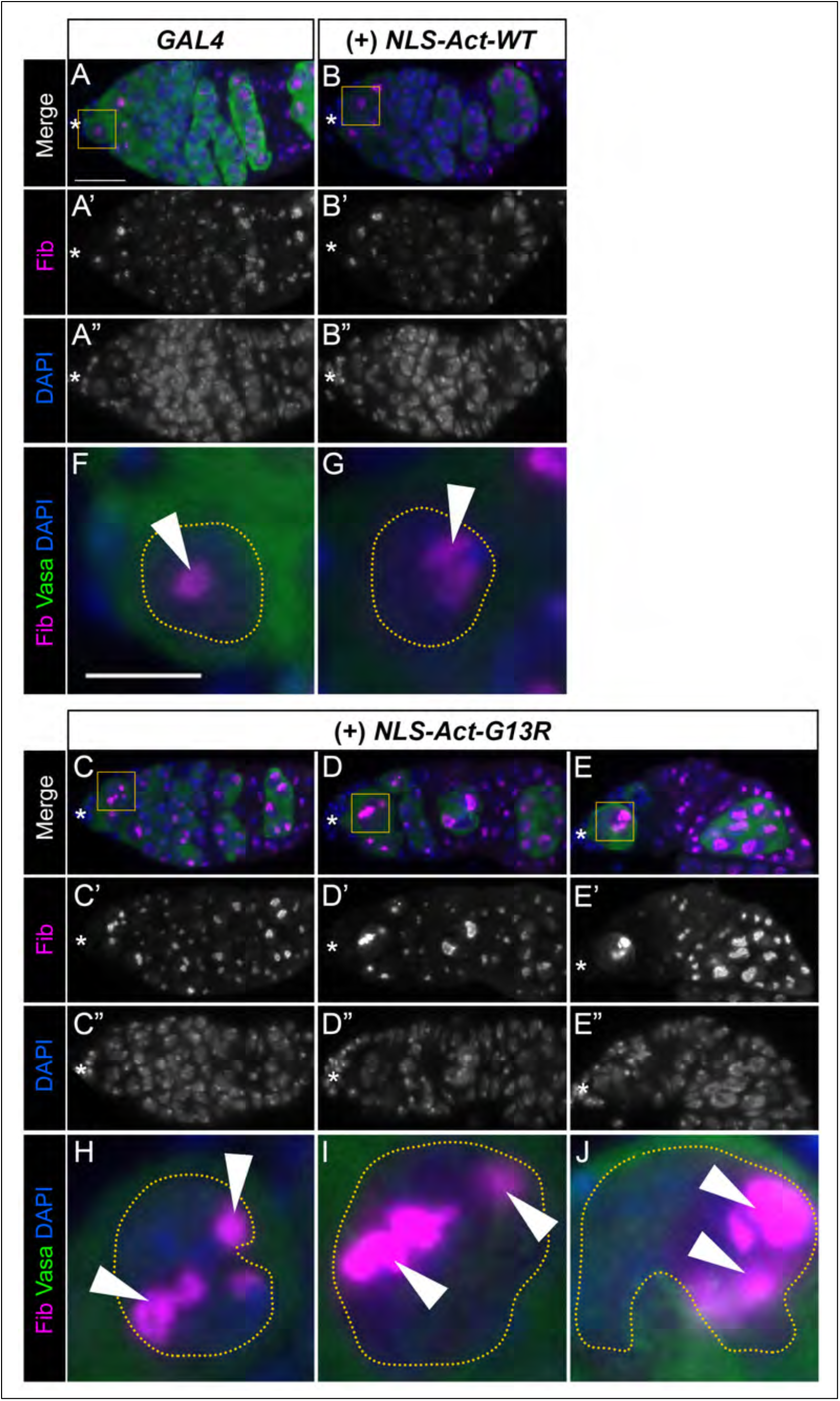
Additional examples of nucleolar morphology changes in control versus NLS-Act overexpression germaria. (A-E”) Maximum projections of 2-5 confocal slices of germaria from 4 day old flies for the GAL4 control (*nos*, A-A”), and germline overexpression of NLS-Act-WT (*nos>NLS-Act-WT*, B-B”), and NLS-Act-G13R (*nos>NLS-Act-G13R*, C-E”) stained for stained for Fibrillarin (Fib, magenta in merge) to mark the nucleolus, DAPI (blue in merge) to mark the nucleus, and Vasa (green in merge) to mark germline cytoplasm. Asterisks designate the anterior tip and GSC niche in each germarium. (F-J) Cropped higher magnification of GSCs in yellow boxed regions of germaria in A-E. Scale bar = 15 μm in A-E”; 5 μm in F-J. Control GAL4 germaria have specific nucleolar (Fib) and nuclear (DAPI) sizes; GSCs have larger nucleoli and nuclei and these size ratios decrease as cells become more differentiated (2-, 4-, 8- and 16-cell cysts) until becoming larger again in regions 2b-3 (A-A”, F). Whereas NLS-Act-WT germaria show the expected nucleolar and nuclear sizes (B-B”, G), NLS-Act-G13R germaria contain germline cells with nucleolar and nuclear hypertrophy which becomes increasingly more severe as germaria lose germline cells (arrowheads, C-E”, H-J). Before nucleolar:nuclear ratios change dramatically, nucleoli begin to change shape and fragment into multiple Fibrillarin puncta (C-C’ and H, yellow arrowheads). As germline cells swell in size, nuclei and nucleoli become more hypertrophied and deform and fragment (D-E” and I-J, arrowheads).

**SFigure 6.**
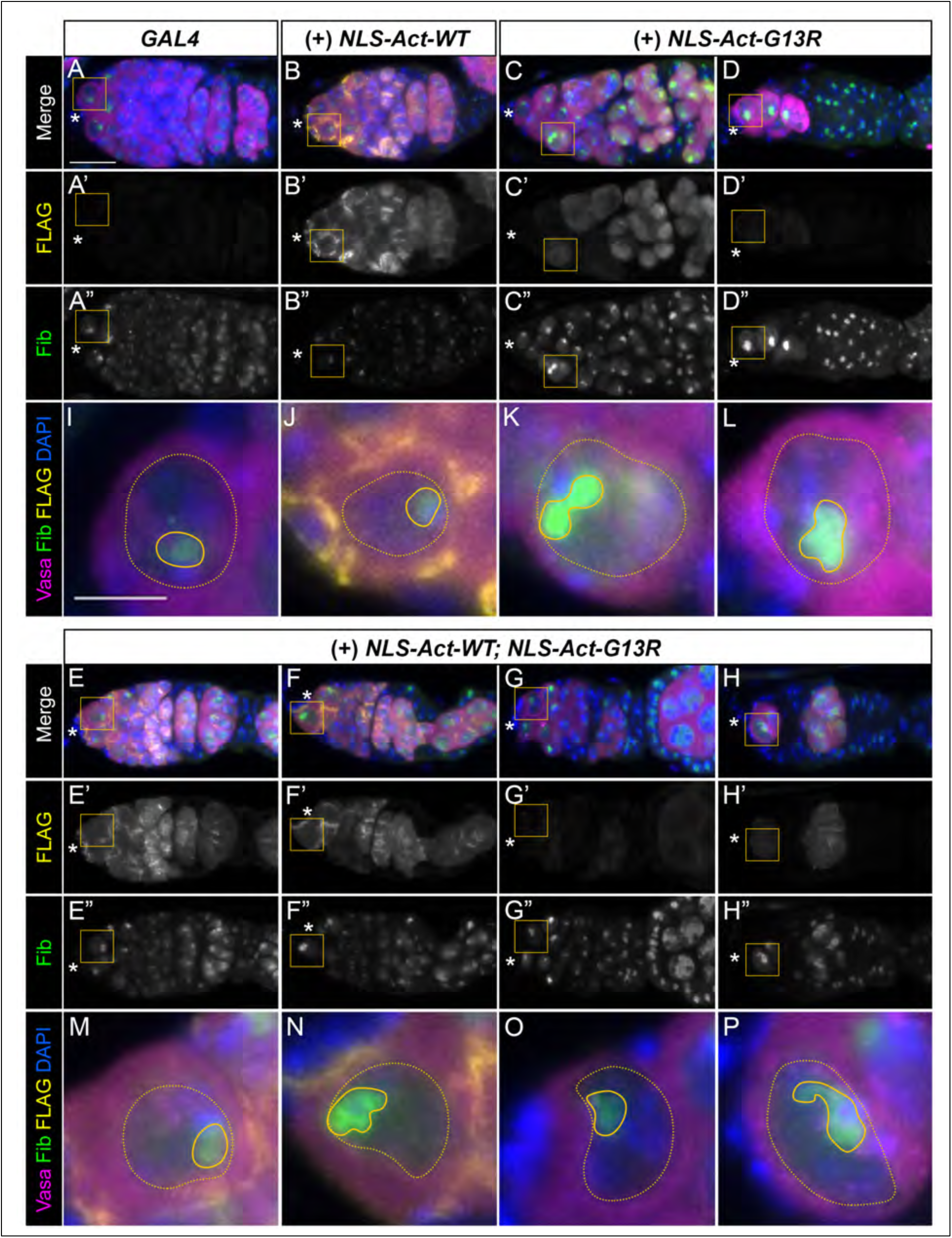
Additional examples of nuclear morphology after restoring nuclear actin polymerization in NLS-Act-G13R overexpressing germaria compared to controls. (A-H’’’) Maximum projections of 2-5 confocal slices of germaria from 4 day old flies for the GAL4 control (*nos*, A-A’’), and germline ovexpression of polymerizable NLS-Act-WT (*nos>NLS-Act-WT*, B-B’’), monomeric NLS-Act-G13R (*nos>NLS-Act-G13R*, C-D’’), and of both polymerizable NLS-Act-WT and monomeric NLS-Act-G13R (*nos>NLS-Act-WT; NLS-Act-G13R*, E-H’’) stained for FLAG (yellow in merge), Fibrillarin to label the nucleoli (Fib, green in merge), Vasa to label the germline (magenta in merge), and DAPI to label the nuclei (blue in merge). Asterisks designate the anterior tip and GSC niche in each germarium. (I-P) Cropped higher magnification of GSCs yellow boxed regions of germaria in A-H. Dotted lines outline the GSC nucleus and solid lines outline nucleoli. Scale bar = 15 μm in A-H”; 5 μm in I-P. FLAG staining is absent in GAL4 controls as expected (A-A’), but exhibits a structured pattern consistent with polymerization when NLS-Act-WT is overexpresed (B-B’). Alternatively, NLS-Act-G13R exhibits a hazy and diffuse FLAG pattern consistent with it blocking polymerization. In both the GAL4 control and NLS-Act-WT overexpression, the germaria contain GSCs and the full range of developing germline cysts (A-B’’). Overexpression of NLS-Act-G13R results in progressively emptying germaria (C-D’’). Co-overexpression of NLS-Act-WT and NLS-Act-G13R restores the presence of structured polymeric actin staining (E-H’’) and rescues germline loss (E-E’’), though some germaria still have an abnormal germline cyst organization (F-F’’) or lose of some germline cells (G-H’’). In GAL4 controls and NLS-Act-WT overexpression, nuclei and nucleoli are larger in GSCs and nucleolar:nuclear size ratios decrease as cells become more differentiated (A-B”). In GSCs, there is a singular round nucleoli (I-J). Inhibiting nuclear actin polymerization (NLS-Act-G13R) disrupts nuclear:nucleolar size ratios across the germarium (C-D”) and causes deformed GSC nucleoli (K-L). Co-overexpressing NLS-Act-WT and NLS-Act-G13R rescues germline loss, and nuclear and nucleolar size and shape of (E-E” and M) is similar to GAL4 controls (A-A” and I). Despite germline cells being restored, co-overexpression of NLS-Act-G13R and NLS-Act-WT is not a perfect rescue, as nuclear hypertrophy is observed (G-H”) and nuclear and nucleolar shape are deformed in some germaria (F-H” and N-P).

**Supplementary Table 1:**
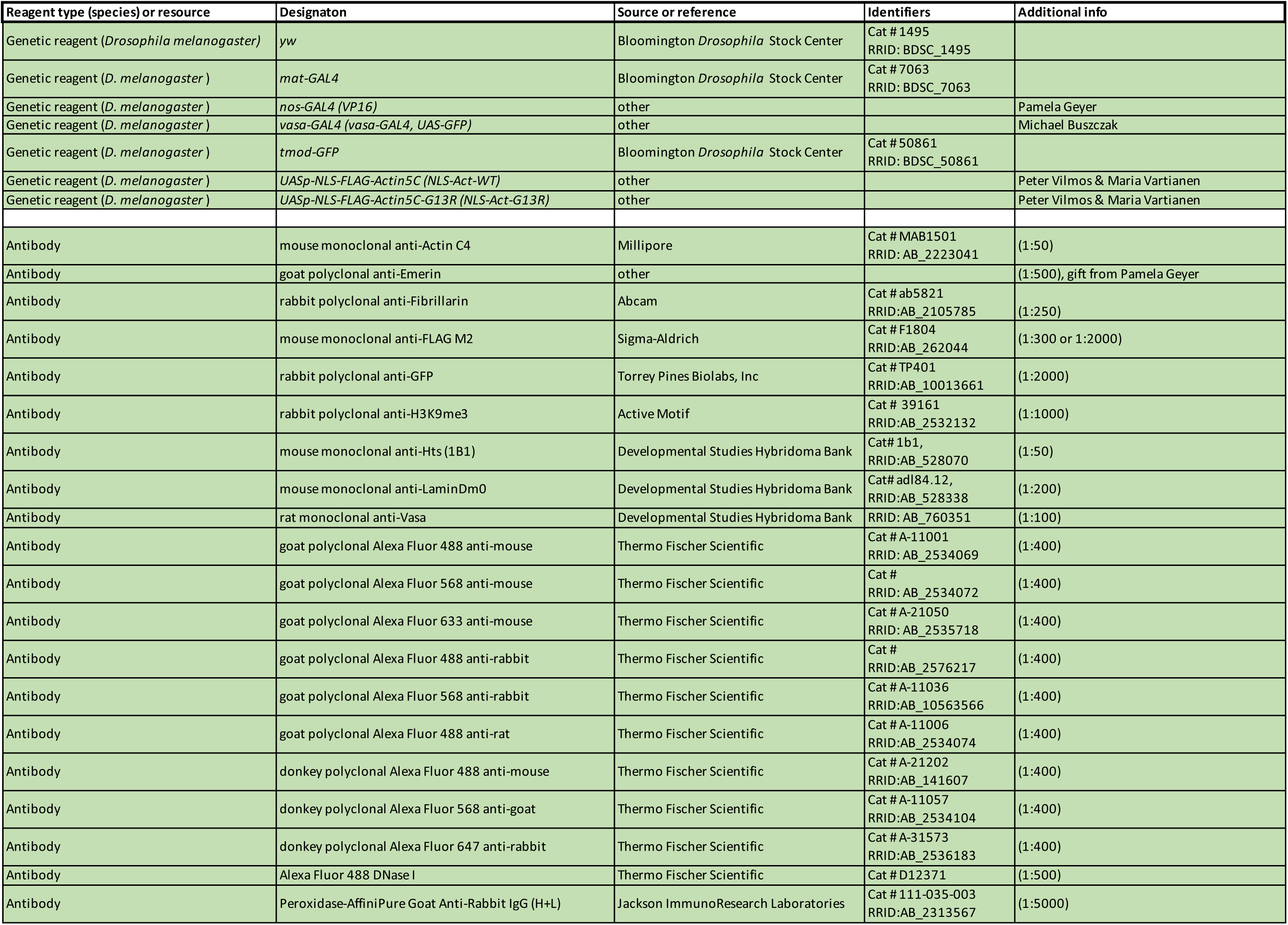
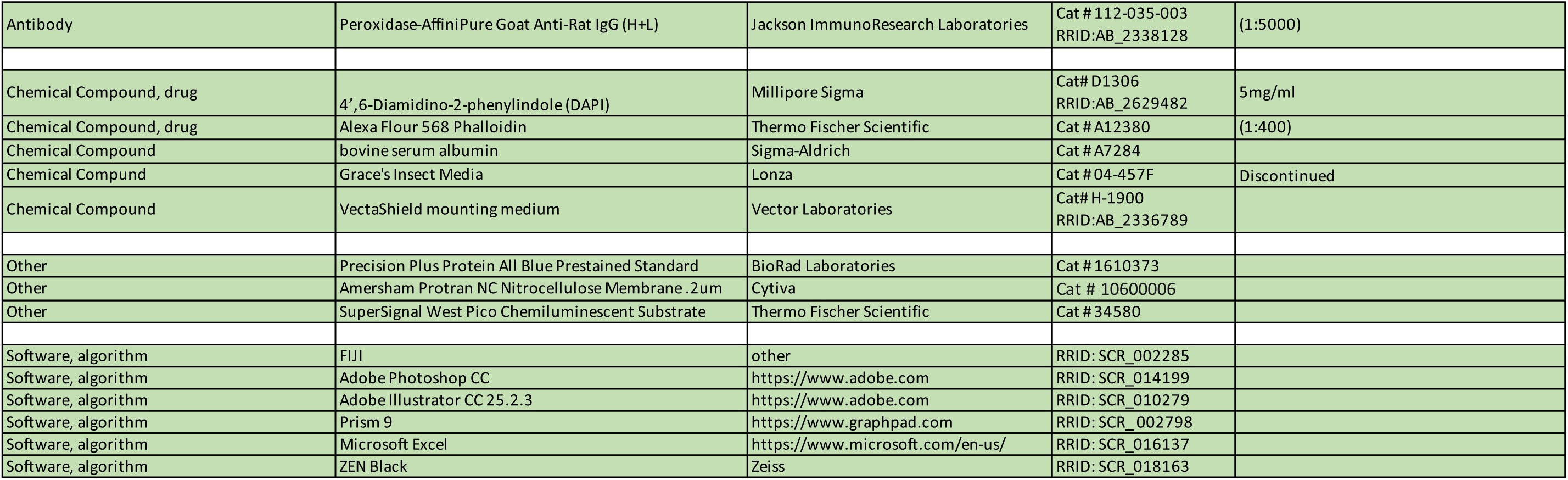
List of reagents used in the study.

